# Root branching in salt requires auxin-independent modulation of LBD16 function

**DOI:** 10.1101/2023.04.25.538210

**Authors:** Yanxia Zhang, Yiyun Li, Thijs de Zeeuw, Kilian Duijts, Dorota Kawa, Jasper Lamers, Kristina S. Munzert, Hongfei Li, Yutao Zou, A. Jessica Meyer, Jinxuan Yan, Francel Verstappen, Yixuan Wang, Tom Gijsberts, Jielin Wang, Nora Gigli-Bisceglia, Timo Engelsdorf, Aalt D.J van Dijk, Christa Testerink

## Abstract

Salinity stress constrains lateral root (LR) growth and severely impacts plant growth. Auxin signaling is indispensable for the regulation of LR formation. Nevertheless, the molecular mechanism of how salinity affects root auxin signaling and whether salt would steer alternative pathway(s) to regulate LR development is unknown. Here we show that the auxin-regulated transcription factor LATERAL ORGAN BOUNDARY DOMAIN (LBD)16, known as an essential player for LR development under control conditions, is regulated by an alternative non-canonical pathway under salinity. Salt represses auxin signaling but in parallel activates an upstream transcriptional activator of LBD16, ZINC FINGER OF ARABIDOPSIS THALIANA 6 (ZAT6). ZAT6 modulates the activity of *LBD16* to contribute to downstream cell wall remodeling, and promotes LR development under salinity stress. Our study thus shows that root developmental plasticity in response to salt stress is achieved by integration of auxin-dependent repressive and salt-activated auxin-independent pathways converging on LBD16 to modulate root branching modulation under salinity.

## Introduction

Plant root system architecture (RSA) acclimation is important for plants’ survival and productivity in response to environmental challenges. In Arabidopsis, lateral roots (LRs) contribute the majority of the mature root system architecture. Their formation and growth are tightly regulated by both internal chemical signals and external environmental cues, for example soil salinity and water availability^1, 2^. LRs are specified within the main root oscillation zone - from the basal meristem zone through the elongation zone- and initiated from xylem pole pericycle (XPP) cells in the differentiation zone^3, 4^. During LR initiation, asymmetric cell division takes place and results in the formation of Stage I lateral root primordia (LRP) in the early differentiation zone, which is followed by early morphogenesis (Stage II through IV) and further meristem organization (Stage V to VIII) of the LRP to traverse the overlaying cell layers and establish emerged LRs^3–5^. Cell wall changes have been observed to occur in the LRP overlaying endodermal cell layer of maize plants as early as in the 1970s^6^. Recently, pectin esterification state at LR initiation sites, as a result of the action of several pectin methylesterases (PMEs) (PME2, 3 and 5) and PME inhibitor 3 (PMEI3), was reported to affect LR formation in Arabidopsis^7^. Additionally, the cell wall-loosening proteins - EXPANSIN (EXP) A1, 14 and 17 - were also reported to promote LR formation^8–10^. Thus, LR development requires coordination of cell division and cell wall modifications to accommodate the emerging LRs.

The plant hormone auxin plays indispensable roles during various stages of LRP development from initiation to LR emergence via different AUXIN RESPONSE FACTOR (ARFs)-mediated transcriptional modules^11, 12^. Arabidopsis ARF7 and ARF19 were shown to mediate LR development by transcriptional regulation of several redundant LATERAL ORGAN BOUNDARY DOMAIN (LBD)-type transcription factors including LBD16, LBD18, LBD29 and LBD33 to mediate LR development^13–15^. Auxin was shown to be upstream of the cell wall remodeling pathway(s) during LR development. For example, auxin induces IDA-HAE/HSL2 ligand-receptor signaling to active a MAPK cascade MKK4/MKK5– MPK3/MPK6 which is required to for the expression of cell wall remodeling genes during LR emergence^16, 17^. The enzymes xyloglucan endotransglucosylases (XTH) 9 and 23, involved in the endotransglucosylation and endohydrolysis of the major cell wall hemicellulose xyloglucan, were shown to act downstream of auxin signaling mediating LR development^16, 18^. In summary, auxin, cell wall modification and their interplays are all playing essential roles in mediating LR development. However, it is not clear how auxin signaling and cell wall modifications during LR development are modulated in response to external environmental signals.

In response to salinity stress, Arabidopsis plants remodel their RSA through modulation of LR emergence and elongation^1, 19–21^, resulting in shorter, fewer LRs in general, although natural variation in the response has been found. Several novel genetic components contributing to the LR developmental plasticity in response to salinity in Arabidopsis have been identified, including *High-affinity K^+^ transporter 1* (*HKT1*) and *CYTOCHROME P450 FAMILY 79 SUBFAMILY B2* (*CYP79B2*)^1^, coding for proteins involved in Na^+^/K^+^ homeostasis and auxin-related metabolite biosynthesis, respectively. Nevertheless, these studies cannot explain how auxin signals are affected during LR development in salt, and how salt impacts the core LR developmental pathways is largely unknown.

Here we report that LBD16 is a crucial mediator of LR developmental plasticity in salt, enabling cell wall remodeling. We show that salt upregulates *LBD16* expression in the main root zones and developing LRP independently of ARF7/19. Simultaneously, we observe a reduction in root auxin levels upon salt stress, revealing the presence of an alternative auxin-independent pathway promoting LBD16 upregulation by salt stress. Yeast-one-hybrid screening combined with transcriptomic analysis and network inference identified additional potential salt stress-induced upstream transcriptional regulators of LBD16. We predicted the C2H2-type transcription factor ZINC FINGER OF ARABIDOPSIS THALIANA 6 (ZAT6), to play a pivotal role in activating *LBD16* in response to salinity and could indeed confirm that ZAT6 binds to the promoter of *LBD16*, acting independent from ARF7/19 to positively regulate *LBD16* expression, contributing to cell wall modifications in roots in response to salinity. Loss-of-function mutants of *ZAT6* displayed salt stress-induced defects in LR development and cell wall composition, similar to the *lbd16-1* mutants. Our study thus provides molecular insights into the coordination of LR developmental plasticity mediated by a salt-activated pathway that positively impacts root branching.

## Results

### *LBD16* is a central regulator of the plasticity of LR development in response to salinity

RSA analyses of the Arabidopsis HapMap natural diversity panel in response to salt stress have identified the previously identified the auxin-dependent transcription factor LBD16 as a candidate locus regulating RSA remodeling in response to salt stress^1^. Since LBD16 had been already known to regulate LR development^13, 14^, we further characterized the RSA of knock-out and complementation lines of *LBD16* under control and salt conditions. We found that the decrease in emerged LR density of two independent *lbd16* mutants in response to salt was more severe compared to the wild type Col-0 (**Fig. 1a, b**, Fig**.S1**). This reduced LR density was not due to main root length differences between Col-0 and the mutants and a genomic region of *LBD16* gene was able to restore the profound decrease in emerged LR density in the *lbd16-1* mutant under both mild (75 mM NaCl) and severe (125 mM NaCl) salt stress conditions (**Fig. S2**). No significant decrease was found in LR density under control conditions in the *lbd16* knock-out mutants compared to Col-0 (**Fig. 1a, b and Fig. S2a, b**), which is in agreement with previous reports showing a functional redundancy of LBD16 with LBD18 and LBD33 under optimal conditions with regard to LR development^13, 22^.

**Fig. 1.**
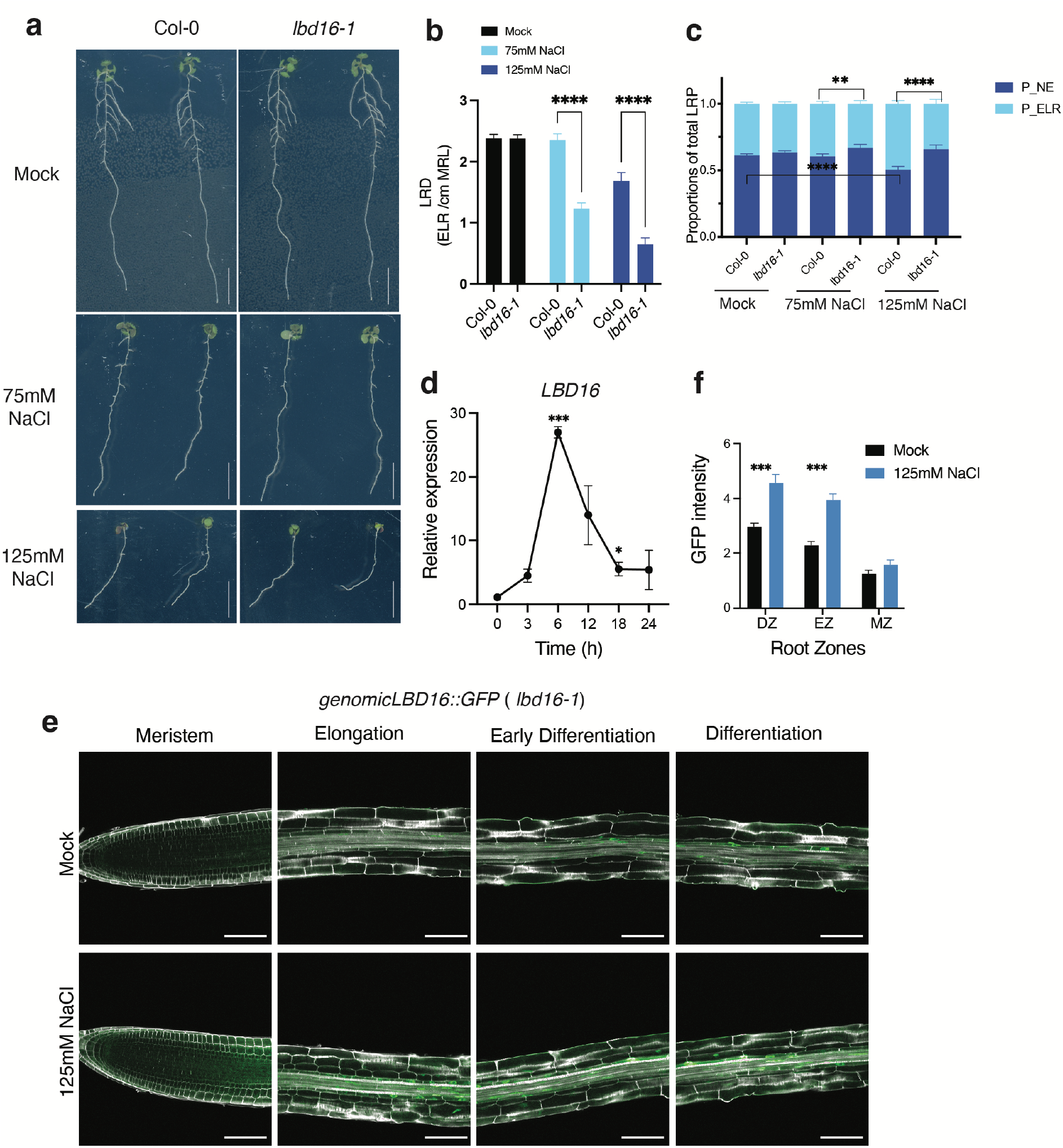
Identification of LBD16 as a mediator for LR development in response to salinity. **a-c**, Phenotypical analysis of emerged LRs (ELR) and non-emerged LRP (NE) in Col-0 and *lbd16-1* in control and salt conditions. **a**, Representative images of root phenotypes of *lbd16-1* compared with Col-0 and *lbd16-1* mutant in control and salt conditions (0 mM, 75 mM NaCl and 125 mM NaCl). Scale bars represent 1cm. **b,** Density of ELR in Col-0 and *lbd16-1* mutant in control and salt conditions (0 mM,75 mM and 125 mM NaCl) (n= 20∼25). **c,** Distribution of ELR and NE LRP in Col-0 and *lbd16-1* mutant in control and salt conditions (0 mM, 75 mM and 125 mM NaCl). Data represents a pool of 4 independent experiments (in each experiment n=7∼13). **d**, Salt-induced *LBD16* expression patterns in 4-week-old hydroponically grown Col-0 wild-type plants (n=4∼5). Relative expression was normalized by both house-keeping gene *At2g43770* and expression at 0 h timepoint. **e,** Expression pattern of genomic *LBD16::GFP* fusion in root zones after treated with 0 mM and 125 mM NaCl for 6h. **f,** Quantification of *gLBD16::GFP* intensities in root zones. Data in **b-d** and **f** represent means ± SEM. Statistical analysis in **b** was done using two-way ANOVA followed by Tukey’s multiple comparison tests. Statistical analysis in **c** was done by fitting a generalized linear model. Statistical analyses in **d** and **f** were done using two-sided Student’s T-test. * P < 0.05, ** 0.05<P<0.01, *** P<0.01, **** P<0.001. Data in a-f represents multiple independent experiments. Scale bars represent 100µm in **e**.

To further understand how *LBD16* may act under saline conditions, we investigated microscopic LRP development in Col-0 and *lbd16-1* in response to mild and severe salt stress. In line with the reported effect of *lbd16* mutant on LR density^13^, we also observed a lower density of non-emerged LRP (Stage I through VII) in *lbd16-1* than in Col-0 under control condition (**Fig. S2c**). Salt stress hampered the LR emergence in Col-0 as more non-emerged primordia were found under 125 mM NaCl (**Fig. 1c**). Compared to Col-0, *lbd16-1* displayed a significantly lower proportion and density of emerged LR under both 75mM and 125 mM NaCl (**Fig. 1c** and **Fig. S2c**), suggesting that LBD16 acts as a positive regulator of the LRP emergence under salt stress.

We further analyzed the dynamic expression patterns of *LBD16* in the roots of four-week-old *Arabidopsis* Col-0 wild-type plants after 0, 3, 6, 12, 18 and 24 h treatment with 125 mM NaCl. *LBD16* expression was enhanced after 3 h and peaked at 6 h after salt treatment (**Fig.1d**). Further investigation of a promoter reporter line expressing *proLBD16::GUS* showed that the promoter activity of *LBD16* was enhanced after 6 h salt treatment with 125 mM NaCl in different regions of the primary root (**Fig. S3a**); from the elongation zone through the differentiation zone, and up to the lateral root zone where lateral roots emerge (**Fig. S3b**). Consistently, an *LBD16genomic ::GFP* (*lbd16-1*) fusion also revealed that *LBD16* expression was enhanced in the main root elongation and differentiation zones by 125 mM NaCl (**Fig. 1e and f**). Salt-induced *LBD16*-expressing cell types were not limited to the root stele but also found in endodermis and cortex layers during LRP development in the differentiation zone and the lateral root zone (**Fig. S3**), implying that LBD16 might play a role during LRP emergence to pass through various root cell layers under salt. Taken together, salt may positively activate an LBD16-mediated pathway in main root zones, which is required for plasticity of LR development.

### Salinity activates an auxin-independent pathway mediating *LBD16* expression

Since *LBD16* was previously shown to be involved in auxin-regulated LR development^14^, we first addressed the question whether salt would affect auxin signaling during LR development. We used a high-resolution C3PO auxin reporter line that contains a three-color reporter by incorporating a DR5v2::n3mTurqoise2 (auxin response) cassette into the construct carrying the R2D2 (auxin input) cassette (pDR5v2:mTurquoise-NLS/pRPS5a:mDII:ntdTomato/pRPS5a:DII:n3xVenus) to visualize auxin dynamics at cellular level during LR development under salt stress^23^. We found that DR5v2 activity was reduced in the root zones including meristem, elongation and early differentiation zones after 6 h treatment with 125 mM NaCl (**Fig. 2a** and **b**), in accordance with decreased IAA levels in Col-0 roots measured by liquid chromatography-tandem mass spectrometry (LC-MS/MS) after 6 h of 125 mM NaCl treatment (**Fig. 2c**). In the lateral root primordia, a significant reduction in the auxin input R2D2 signal (indicated by mDII/DII) was also observed in Stage I and II after 6 h 125 mM NaCl treatment, while we did not detect significant changes in the auxin output signal (DR5v2) in developing early stage LRP in response to salt (**Fig. S4a** and **b**). Although these results are in line with salt inhibiting LR formation (**Fig. 1c** and **Fig. Sc**), they do raise the question how salt enhances *LBD16* expression in roots (**Fig. 1e** and **f**) and whether this could occur independent of auxin.

**Fig. 2.**
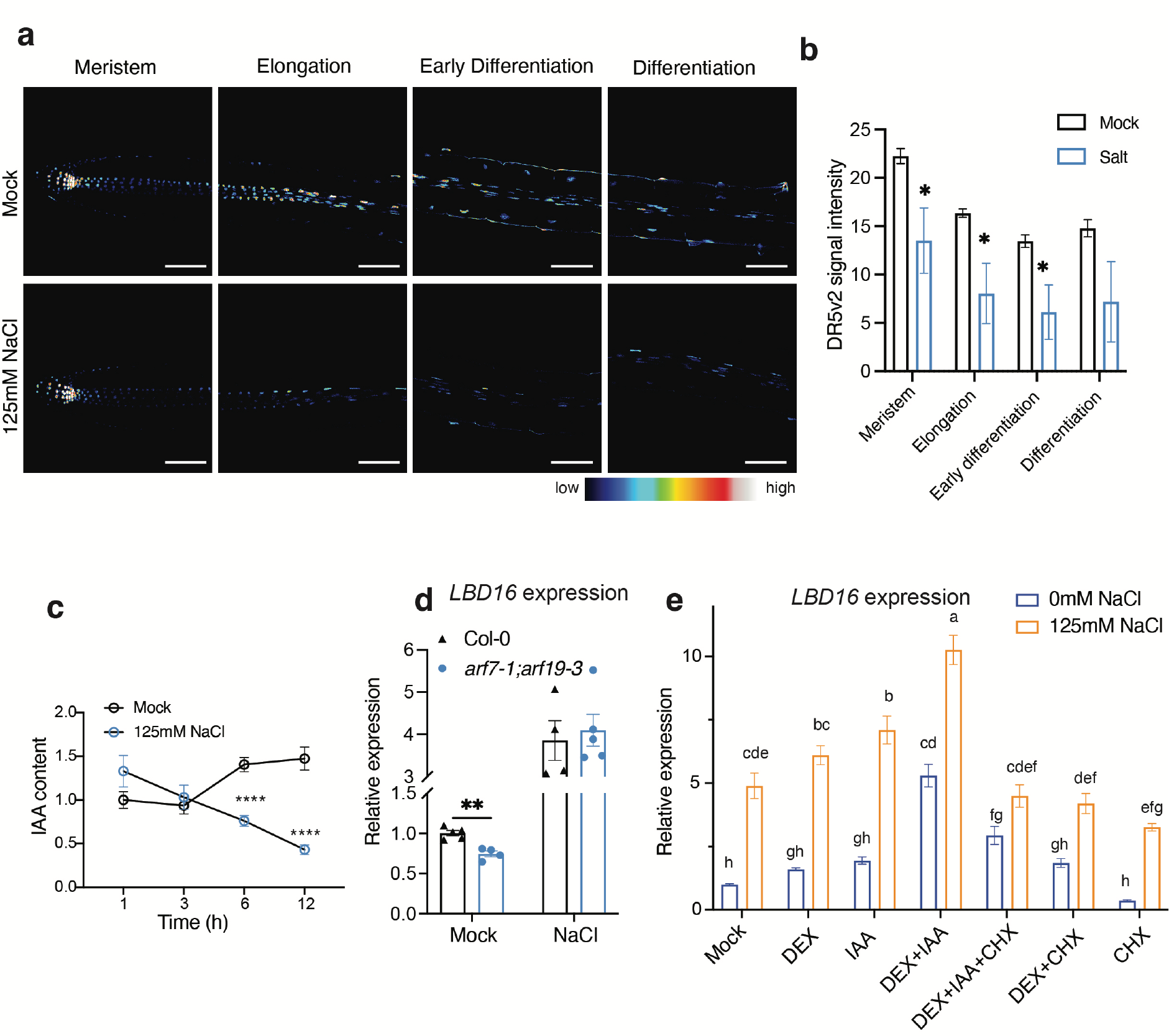
Salinity activates an auxin-independent pathway mediating *LBD16* expression. **a,** auxin output signals indicated by DR5v2 in main root zones. **b,** Quantification of DR5v2 signal intensity in root zones including meristem, elongation zone, and (early) differentiation zone. **c,** Relative IAA content normalized by internal standard in the roots of 7-day-old Col-0 seedlings by LC-MS/MS after 1, 3, 6 and 12 h 125 mM NaCl treatment (n=4). **d,** Relative expression of *LBD16* compared with Col-0 under control condition in the roots of 5-day-old seedling of *arf7-1;arf19-3* double mutants and Col-0 plants after 6 h 130 mM NaCl treatment (n=4∼5). **e,** Relative expression of *LBD16* in the *pARF7::ARF7::GR* (in *arf7-1;arf19-1*) plants were treated with 1 µM *IAA*, 2 µM dexamethasone (DEX) and/or 10 µM cycloheximide (CHX) inducible lines in control and salt conditions after 6h 125 mM NaCl treatment in comparison with Mock treatment. Expression values were normalized by house-keeping gene *At2g43770* in c and d. Statistical analyses in **b** and **c** were done using two-sided Student’s T-test. Statistical analyses in **d and e** were done using two-way ANOVA, respectively. * P < 0.05, ** 0.05<P<0.01.

As *LBD16* expression in control conditions is induced by ARF7 and ARF19 acting downstream of auxin, we evaluated *LBD16* expression in the *arf7-1;arf19-3* double knock-out mutant. Interestingly, although under control conditions *LBD16* transcripts were decreased in the *arf7-1;arf19-3* double mutant as compared with Col-0, after 6 h salt treatment *LBD16* expression was increased in the *arf7-1;arf19-3* double mutant compared to Col-0 (**Fig.2d**). This result implies that *LBD16* expression is regulated by an ARF7/ARF19 independent pathway under salt conditions (**Fig.2d**). Since ARF7 was reported to be able to bind to the promoter region of *LBD16* and regulate its expression^14^, to further investigate the regulation of *LBD16* by ARF7 under salt stress, we made use of a *ProARF7:ARF7-GR/arf7-1;arf19-1* transgenic line with dexamethasone (DEX)-inducible ARF7 activity^14^. In control conditions, in line with a previous study^14^, *LBD16* expression was induced by the treatment with DEX plus IAA in the roots of *ProARF7:ARF7-GR/arf7-1;arf19-1* seedlings, and this high expression was maintained by the combination of IAA and DEX in the presence of the protein synthesis inhibitor cycloheximide (CHX) to inhibit new protein synthesis of ARF7 (**Fig.2e**), suggesting LBD16 is a primary target of ARF7 under control conditions. In response to salt stress, *LBD16* expression was induced significantly by the combination of DEX and IAA in the presence of CHX already early after 3 h treatment when compared with treatments with DEX and IAA alone (**Fig.S4c**), indicating that LBD16 is primarily regulated by auxin-ARF7 signaling during early salt response at 3 h. However, we observed that the presence of CHX together with DEX and IAA to induce ARF7 activity was not able to promote *LBD16* expression after 6 h salt treatment, and *LBD16* expression was significantly enhanced by salt in the roots of *ProARF7:ARF7-GR/arf7-1;arf19-1* seedlings even without ARF7 activity under mock treatment in the presence of CHX alone (**Fig. 2e**). These data support our hypothesis that additional player(s) would regulate *LBD16* expression in response to a prolonged salt stress, besides ARF7 and ARF19.

### Identification of novel upstream regulators of *LBD16* acting in response to salt stress

To expand our understanding of the molecular mechanism underlying the salt-induced LR plasticity phenotype of *lbd16* mutants and the observed salt-induced, auxin-independent regulation of *LBD16* expression, we set out to identify possible additional upstream regulator(s) of *LBD16* in response to salt stress. To this end, we performed yeast one hybrid (Y1H) screening by using a 1309- bp promoter region upstream of the *LBD16* start codon as a bait, and a collection of 1956 Arabidopsis transcription factors as prey to survey the putative upstream regulators of *LBD16*^24^. As a result, more than 300 transcription factors were identified to bind to the promoter region of *LBD16* with high presentation of bHLH, AP2-EREBP, C2H2 and MYB -type transcription factors (**Supplementary dataset 1** and Fig. **S5a**). To further prioritize candidate transcription factors upstream of *LBD16* involved specifically in LR development in response to salt stress, we carried out a comparative analysis of public transcriptome datasets on root cell layers in response to salt stress^25^ and gene expression during LR development^26^. We found that 6499 out of 9193 salt-induced differentially expressed genes (DEGs) in Arabidopsis root cell layers overlapped with genes induced during LR development (9581 in total) (**Fig.3a**). We then compared these DEGs with putative LBD16 upstream regulators including 91 candidates from our Y1H screening, and ARF7 and ARF19, which resulted in a subset of 94 transcription factors including LBD16 itself that may be involved in LR development in response to salt stress (**Fig.3a** and **Supplementary dataset 2**).

We next performed a network inference analysis to construct an *LBD16*-associated network using GRNBoost2^27^ based on the salt-induced time-series co-expression data of these 94 genes in the root stele cells from the salt response dataset^25^(**Supplementary dataset 3**). To test the robustness of the network prediction, we computed the network inference 100 times and ranked all the predicted direct regulators of *LBD16* by how frequent they were predicted as an *LBD16* upstream direct regulator. We also performed the same analysis using the expression data under control conditions (**Supplementary dataset 3**). As a result, we were able to identify 10 transcription factors that may regulate *LBD16* expression in control conditions, and predicted 13 to act specifically in salt conditions (**Fig.3b**). Among the potential salt-specific LBD16 upstream regulators, 5 of them were predicted with 100% frequency, including AKS2(AT1G05805), ANAC001(AT1G01010), ZAT6(AT5G04340), BHLH007(AT1G03040) and UNE12(AT4G02590) (**Fig. 3b** and **Supplementary dataset 3**).

**Fig. 3.**
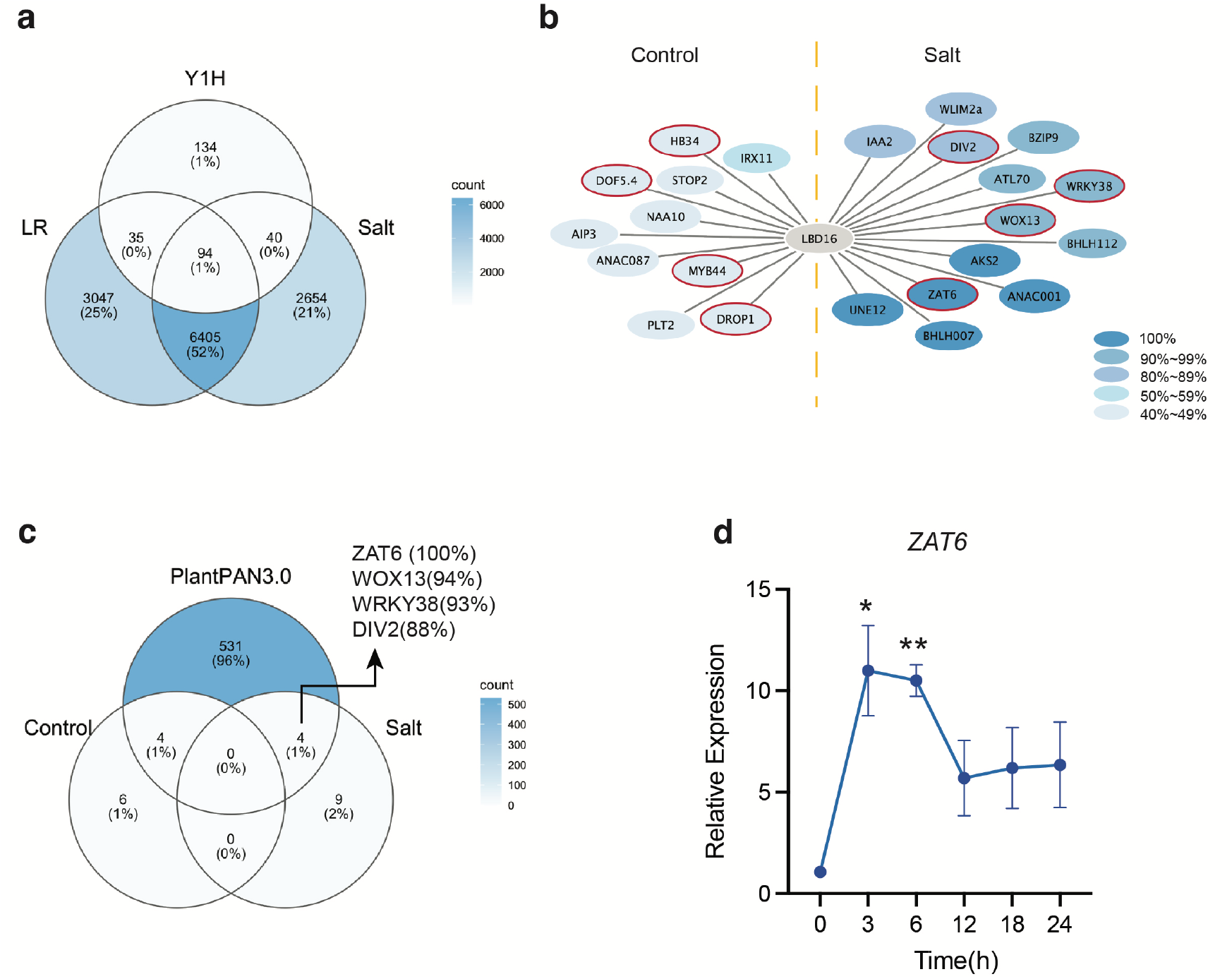
Identification of novel upstream regulators of LBD16 in response to salt stress. **a,** novel upstream regulators of *LBD16* mediating LR development in response to salt stress were selected by a comparative analysis among salt-induced root cell-types specific transcriptomic response^25^, LR development-associated genes^26^ and putative *LBD16* upstream regulators via yeast one-hybrid screening among 1956 *Arabidopsis* transcription factors^24^. **b**, Network inference prediction of LBD16 upstream direct regulators by GRNBoost2 under control and salt conditions. The topology of the network shows genes that could directly regulate *LBD16*. Color (blue) intensity indicates the robustness of the network prediction (100% to 40%). **c**, Comparative analysis of the *LBD16*-associated upstream network with *LBD16* promoter analysis on the basis of ChIP-seq evidences in PLANTPAN3.0^28^. **d,** Salt-induced expression pattern of one of the novel upstream regulators *ZAT6*. Data were collected using roots of 4- week-old hydroponically grown Col-0 plants normalized by house-keeping gene *At2g43770* and expression of 0 h time point (n=4). Statistical analysis in **d** was done using two-sided Student’s T-test. * P < 0.05, ** 0.05<P<0.01.

To narrow down the salt-induced candidates for further functional characterization, we performed an *LBD16* promoter analysis using the PlantPAN3.0 database to screen for upstream transcription factors that were shown to bind the *LBD16* promoter directly in existing ChIPseq experiments^28^. By comparative analysis of both the salt and control networks with the upstream regulator candidates from PlantPAN3.0 (**Supplementary dataset 3**), we identified 4 candidate salt-specific transcription factors that can directly bind the *LBD16* promoter; ZAT6 (AT5G04340), WOX13(AT4G35550), WRKY38 (AT5G22570), and DIV2 (AT5G04760) (**Fig. 3c**). Among these only ZAT6 was predicted with 100% frequency in our network analysis (**Fig.3c and Supplementary dataset 3**). Further gene expression analysis showed that *ZAT6* expression peaked at 3 h after 125 mM NaCl treatment prior to the salt-induced expression peak of *LBD16* (**Fig. 1d and Fig.3d**), suggesting that *ZAT6* expression may be up-regulated as early as 3 h after salt exposure to activate *LBD16* transcriptional activity at 6 h of salt treatment. To examine whether ZAT6 is part of the *ARF7/19-LBD16* path, we also analyzed *ZAT6* expression in the roots of the *arf7-1;arf19-3* double mutant and the *ProARF7:ARF7-GR/arf7-1;arf19-1* line, and found that *ZAT6* expression was not affected by presence of ARF7 and ARF19 in either control or salt conditions (**Fig. S5b** and **S5c**). Collectively, our results reveal that ZAT6 is a potential novel upstream player regulating *LBD16* expression in response to salinity in parallel with, but acting independently of ARF7 or ARF19.

### ZAT6 acts as a positive upstream regulator of *LBD16* in response to salt stress

To understand the transcriptional regulation of *LBD16* by ZAT6, we further verified the interaction of ZAT6 with the *LBD16* promoter using dual-luciferase assays in *Nicotiana Benthamiana* leaves. We fused the 1309-bp region of the *LBD16* promoter to the FIREFLY LUCIFERASE (LUC) to generate the reporter construct *pLBD16::LUC*. The *ZAT6* coding region driven by the cauliflower mosaic virus (CaMV) 35S promoter was used as an effector construct. Both the reporter and effector constructs together with the internal control RENILLA LUCIFERASE (REN) driven by the 35S promoter were co-infiltrated into *N. benthamiana* leaves using the *A. tumefaciens* AgL0 strain. Notably, co-infiltration of *35S::ZAT6* with *pLBD16::LUC* resulted in a higher relative luciferase activity compared with an empty 35S vector control (**Fig. 4a**). To understand whether ZAT6 is acting upstream of *LBD16* under salt stress in *planta*, we assessed the expression of *LBD16* in roots of the loss-of-function mutants *zat6-1* (SALK_061991C) and *zat6-2* (SALK_050196) as well as a constitutive repression line of ZAT6 by using a SUPERMAN REPRESSIVE DOMAIN X (SRDX) fusion, namely *35S::ZAT6::SRDX*. In both *zat6* mutant alleles (**Fig. S6a)**, a significant decrease in *LBD16* expression was observed after 24 h of 125 mM NaCl treatment, while *LBD16* expression in control conditions was not affected (**Fig.4b**). Similarly, *LBD16* expression in the *35S::ZAT6::SRDX* line was significantly lower compared to Col-0 after 24h salt treatment (**Fig.4c**). Together these results identify ZAT6 as a positive upstream regulator of *LBD16*, which is required for the salt-induced increase in *LBD16* expression in roots.

**Fig. 4.**
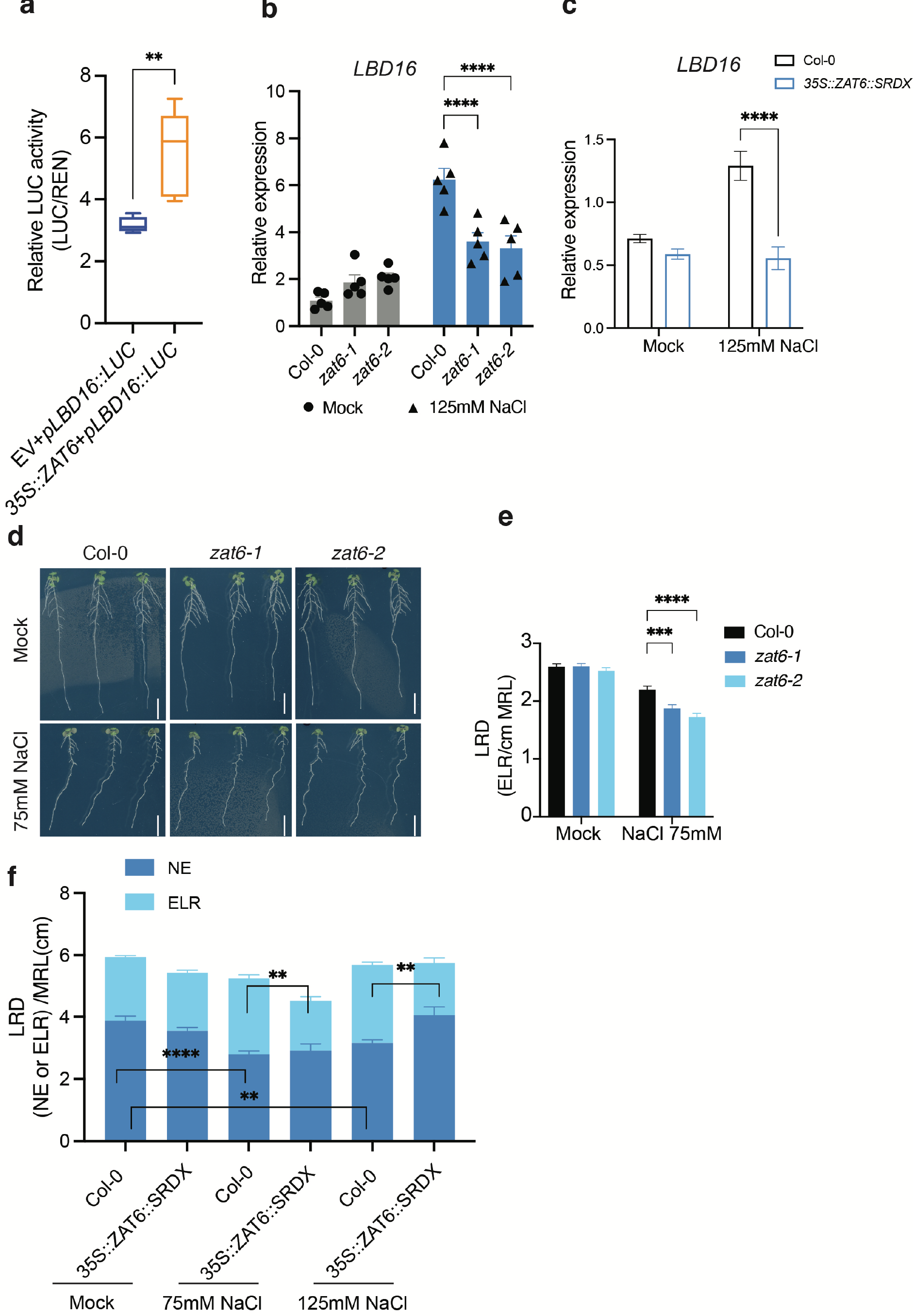
ZAT6 acts upstream of LBD16 in response to salt stress. **a,** Overexpression of *ZAT6* (*35S::ZAT6*) enhanced LUC expression driven by *LBD16* promoter in *Nicotiana Benthamian* leaves (n=5). **b,** Relative expression of *LBD16* in response to 125 mM NaCl treatment in Col-0, *zat6-1* and *zat6-2* mutant alleles quantified by RT-qPCR. **c**, Expression of *LBD16* in roots of *35S::ZAT6::SRDX* line after 24 h treatment by mock and salt (125 mM NaCl) conditions (n=4∼5 pools of 60 roots). **d**, Representative images of root phenotypes of Col-0, *zat6-1* and *zat6-2* mutants under mock and 75 mM NaCl. Scale bars represent 1cm. **e,** Emerged LR density of *zat6-1* and *zat6-2* mutants compared with Col-0 under mock and 75 mM NaCl. Data were collected from 3 independent experiments (3 pools of 20∼25 roots). **f,** Density of emerged LRs and non-emerged LRP in Col-0 and *35S::ZAT6::SRDX* line under mock, 75 mM NaCl and 125 mM NaCl conditions (n=10∼11). Data represent results of two independent experiments (n=10∼13). Data in **a**, **b**, **c**, **e** and f represent means ± SEM. Expression data in **b** and **c** were obtained from 4 biological replicates (approx. 40∼45 roots were pooled as one replicate) and presented as the relative expression to mock condition after normalization by the reference gene *At2g43770*. Root RNA samples were obtained from 7-day old seedlings transferred to 0.5x MS medium supplemented with 0 or 125 mM NaCl for 24 h. Statistical analyses in a and b were done using two-sided Student’s T-test. Statistical analyses in **c, e** and **f** were done using two-way ANOVA, followed by Tukey’s multiple comparison tests. * P < 0.05, ** 0.05<P<0.01, *** P<0.01, **** P<0.001.

To further investigate whether *ZAT6* is involved in LR formation under salinity, we transferred 4-day-old seedlings of Col-0, *zat6-1* and *zat6-2* lines to agar plates containing 0 mM or 75 mM NaCl for 6 days and screened their RSA traits (**Fig. 4d** and **e**). No difference was found in main root length or average lateral root length under either control or salt stress conditions between Col-0 and the mutant plants (**Fig.4d and Fig. S6b, c**). Yet, a reduced emerged LR density was observed in *zat6-1* and *zat6-2* mutants compared with Col-0 under 75 mM NaCl treatment (**Fig.4d, e**). The LR density reduction was evidently due to the reduction in the number of emerged LRs under 75 mM NaCl (**Fig.S6d**), as the main root length did not change significantly (**Fig. S6b**). We further evaluated LR development by scoring the emerged LRs and non-emerged LRP in *35S::ZAT6::SRDX* line under both control and salt conditions. We observed that the density of emerged LRs in the *35S::ZAT6::SRDX* line was significantly decreased compared with Col-0 under 75 mM and 125 mM NaCl treatments whereas there was no difference in main root length between these two genotypes **(Fig.4f** and **Fig.S7**), consistent with the significantly lower *LBD16* expression in the *35S::ZAT6::SRDX* line compared with Col-0 in response to salt stress (**Fig. 4c**). Similar to the *lbd16-1* mutant (**Fig.S2c)**, non-emerged LR density was not reduced in the *35S::ZAT6::SRDX* line in response to salt stress while salt suppressed the formation of LR in Col-0 (**Fig. 4f**). Collectively, the data suggest that *ZAT6* is required for LRP development and LR emergence under salinity stress conditions.

### The *ZAT6-LBD16* pathway regulates cell wall remodeling during salt stress

To gain insight into the molecular mechanisms of *LBD16-*mediated LR developmental plasticity in response to salinity, we analyzed transcriptomes of roots of 8-day-old Col-0 and *lbd16-1* mutant seedlings treated with 130 mM NaCl for 6 h by RNA sequencing (RNA-seq). Further quality control analysis using principal component analysis (PCA) and hierarchical clustering methods presented distinct expression patterns between salt and control conditions (**Fig. S8a-c**). In total, 384 genes were differentially expressed (FDR:<0.05) between *lbd16-1* and Col-0 in control conditions, while 1254 significant DEGs were observed under salt treatment. Among these 1254 genes, 101 genes overlapped with DEGs in control condition and 1153 DEGs were found specifically for salt stress (**Fig.5a** and **Supplementary dataset 4**). To identify key molecular processes involved in the salt-induced *LBD16*-mediated root branching plasticity, we further performed gene ontology (GO) analyses for salt-specific DEGs (**Supplementary dataset 4**). Strikingly, some of the most significantly overrepresented GO terms we found were linked to cell wall organization (biological process, BP), cell wall (cellular compartment, CC) and xyloglucosyl transferase activity (molecular function, MF) (**Fig. 5b, Fig. S8d and Supplementary dataset 4**). Further investigation of the expression profiles of the genes enriched in these cell-wall related GO terms suggested that salt stress more strongly down-regulated the expression of pectin- and xyloglucan-related genes in the *lbd16-1* mutant compared with Col-0, while a subtle difference in their expression profiles was observed between *lbd16-1* and Col-0 in control conditions (**Fig. 5c)**. Among them, for example, *PME2*, *GAUT7*, *XTH19* were previously shown to be expressed in developing LRs^7, 29^. Consistently, salt stress significantly enhanced the expression of *PME2* and *XTH19* in Col-0, whereas the induction was abolished in the *35S::ZAT6::SRDX* line likely due to the repressed activity of *LBD16* by ZAT6 (**Fig. 4c and Fig. S9**). Together, our data suggest that the ZAT6-*LBD16* regulon is involved in salt-induced cell wall remodeling in response to salinity in roots.

**Fig. 5.**
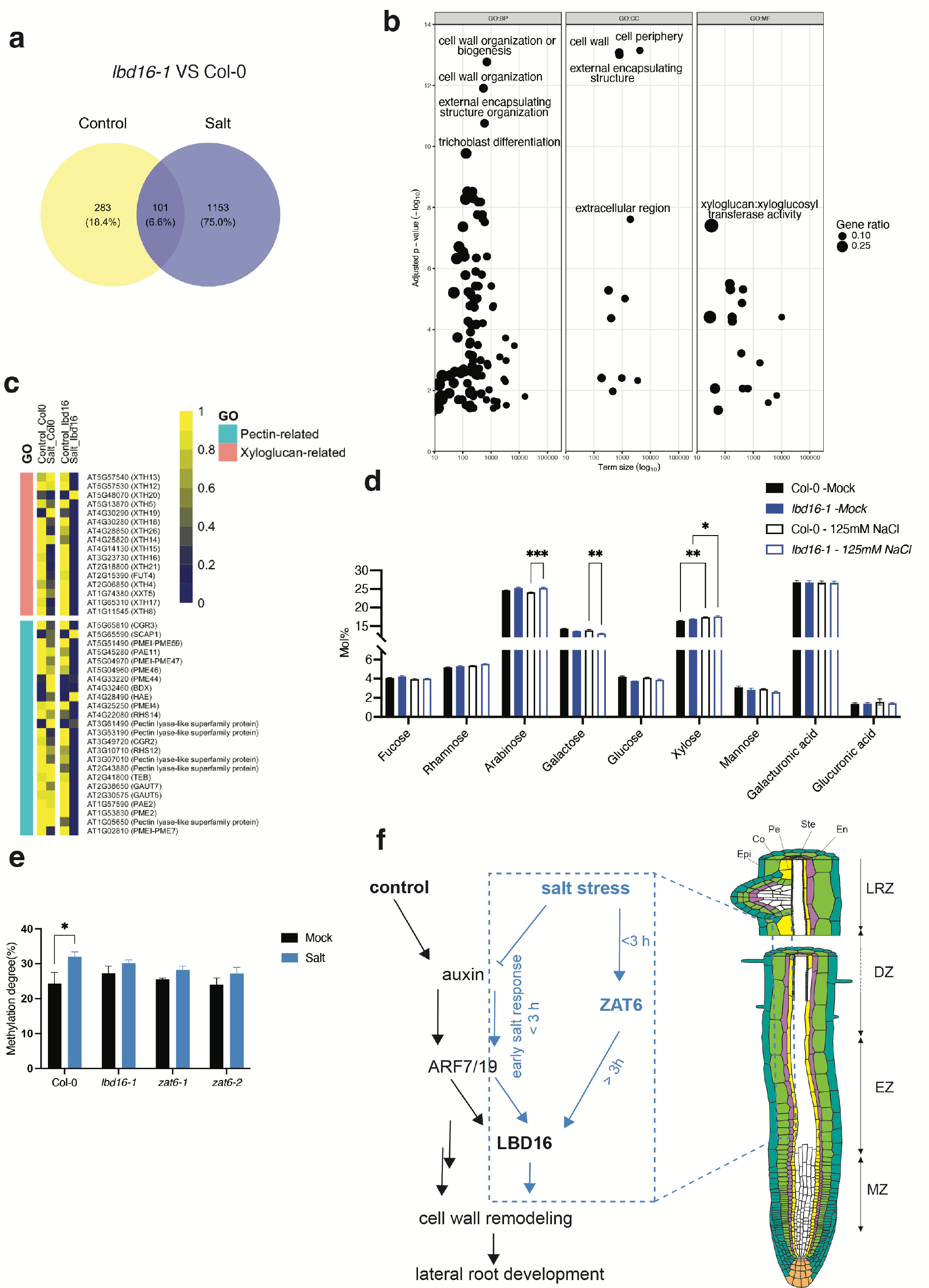
The *ZAT6-LBD16* path regulates cell wall remodeling during salt stress. **a,** Venn diagram representing the number of differentially expressed genes (DEGs) comparing 8-day-old *lbd16-1* mutant with Col-0 after treatment with 0 mM and 130 mM NaCl for 6 h (no cutoff for log-fold changes, P <0.05). **b,** GO term analysis of the 1153 salt-induced genes in **a**. BP, biological process; CC, cellular compartment; MF, molecular function. Y-axis indicates the significance of the GO terms. X-axis indicates term size. **c**, Heatmaps showing the relative expression of genes listed in pectin- and xyloglucan-related GO terms in the roots of Col-0 and *lbd16-1* in control and salt conditions. **d,** Cell wall monosaccharides analysis of roots of 7-day old *lbd16-1* and Col-0 after transferring to agar plates containing 0 mM or 125 mM NaCl for 24 h. Data are mean ± SEM. Data were obtained from 4 independent biological replicates containing 60 ∼80 roots. Statistical analysis was done using two-way ANOVA, followed by Tukey’s multiple comparison tests. * P < 0.05, ** 0.05<P<0.01, *** P<0.01. **e,** The degree of pectin methylation in Col-0, *lbd16-1*, *zat6-1* and *zat6-2* lines under mock and 125 mM NaCl conditions. Data were collected from roots of 7-day old seedlings after transferring to 0 mM or 125 mM NaCl for 72 h (n=4 pools of 60 ∼80 roots). **f,** Schematic diagram of LBD16-mediated molecular network in the main zones guiding lateral root development under control (indicated by black arrows) and salt stress conditions (indicated by blue arrows). The blue dashed boxes highlight the molecular pathways identified by this study. Under control condition, auxin and its downstream response factor ARF7 was reported to be involved in cell wall remodeling process during lateral root development^16^, in which the role of LBD16 was unclear. During early salt stress (< 3h) response, root auxin signaling is inhibited while auxin signaling is still contributing to the regulation of LBD16. Meanwhile, salt activates ZAT6 which is further regulating LBD16 during long-term (> 3h) salt stress response to mediate downstream cell wall remodeling and subsequently lateral root development. Various root cell layers were presented by different colors. White, yellow, purple, light green and dark green represent stele (Ste), pericycle (Pe), endodermis (En), cortex (Co) and epidermis (Epi), respectively. MZ, EZ, DZ and LRZ denote meristematic zone, elongation zone, differentiation zone and lateral root zone (root zone with emerged LRs), respectively.

To assess how the salt-induced *ZAT6-LBD16* pathway affects downstream cell wall modification, we surveyed the cell wall monosaccharide composition changes in the roots of Col-0, *lbd16-1*, *zat6-1* and *zat6-2* mutant lines in control conditions and after 24 h of 125 mM NaCl treatment by high-performance anion-exchange chromatography with pulsed amperometric detection (HPAEC-PAD) of isolated alcohol-insoluble residue cell wall material (AIR). Salt exposure induced differences between Col-0 and *lbd16-1* in the relative amounts of arabinose and galactose, while the relative xylose amount was increased in both lines after salt treatment (**Fig. 5d**). Even if the overall salt-induced cell wall monosaccharide composition in *zat6-1* and *zat6-2* relative to Col-0 was similar to that of the *lbd16-1* mutant, we did not observe significant changes in the cell wall matrix composition of the *zat6* mutants after 24 h or 72 h salt treatment (**Fig. S10**). Since we found salt-induced changes in *PME* and *PMEI* gene expression in the *lbd16-1* mutant relative to Col-0 (**Fig. 5c**), which were reported earlier to be highly relevant for LR development^7^, we further assessed the degree of pectin methylation in the roots of Col-0, *lbd16-1*, *zat6-1* and *zat6-2* mutant lines. We detected a significantly higher degree of pectin methylation in Col-0 after 72 h salt treatment, while the changes in *lbd16-1*, *zat6-1* and *zat6-2* mutant lines were much less noticeable (**Fig. 5e**). This is in agreement with the fact that salt enhanced *PME2* expression in Col-0 whereas it is not altered by salt in the *35S::ZAT6::SRDX* line, suggesting that PME2 expression is dependent on ZAT6 activity and that salt-induced regulation of pectin methylation requires LBD16 and ZAT6 (**Fig. S9**). In summary, these data indicate that salt stress activates the *ZAT6-LBD16* module to regulate downstream cell wall modification, positively contributing to LR developmental plasticity under salinity.

## Discussion

Plant lateral roots play important roles in acquisition of water and nutrients. Flexibility in when and where to branch is essential for a plant to help reach optimal RSA for a dynamic soil environment. We report here that *LBD16* plays a major role in mediating root branching plasticity in response to salinity. We identified ZAT6 as a positive regulator that functions independently of auxin-induced ARF7/19 signaling to induce *LBD16* expression in long-term salt-stressed roots. Salt attenuates root auxin signaling since early stress response, whereas it enhances the ZAT6-LBD16 module to facilitate downstream cell wall remodeling, likely to promote LR development in response to salinity stress, bypassing the canonical root branching pathways that function in control conditions (**Fig.5f**).

Auxin plays an essential role during LR formation and its downstream transcription factor LBD16 was demonstrated to be required for LR initiation occurring in the early differentiation zone of the *Arabidopsis* root^3, 13^. Using the auxin reporter line C3PO, we provide evidence of salt-suppressed auxin signaling (DR5v2 signals) in the main root zones where LRs are primed (oscillation zone) and initiate (differentiation zone), and reduced auxin input signal (mDII/DII) during the early stages of LRP development (Stage I through Stage IV) upon salt stress (**Fig.2a- b, Fig.5a**). These data were further corroborated by the quantification of IAA content in salt-stressed roots (**Fig. 2a**). The reduction in auxin signaling may contribute to the decreased density of non-emerged LRP in Col-0 (**Fig. S2c**). However, salt did not affect the density of non-emerged LRP in *lbd16-1* mutants (**Fig. S2c**), suggesting a positive role for LBD16 during LR formation under salt conditions. In agreement with this, our data reveal that salt halts LRP emergence leading to a decrease in emerged LR density in *lbd16-1* relative to Col-0 (**Fig. 1a-c and Fig.S2c**). Moreover, salt-enhanced *LBD16* promoter activity and expression were not only detected in the differentiation zone but also in the elongation zone (**Fig. 1e and Fig. 3a**), suggesting LBD16 may have additional role(s) in the developmental processes prior to LR initiation during salt stress, such as LR priming^3^. Due to reduced auxin signaling upon salt stress, it is not likely auxin is the trigger for salt-induced *LBD16* expression (**Fig. 2a-c**). Together, it appears that salt stress has an overall negative effect on auxin accumulation and signaling in the main root zones involved in both LRP development and emergence, whereas it also activates positively acting pathway(s) in which LBD16 is required to promote LRP development under salinity. In the *lbd16* mutants, this positive contribution is lacking, resulting in the observed reduced root branching phenotype.

Our network inference analysis indicates that, besides ARF7 and ARF19, various transcription factors, such as ZAT6 (AT5G04340), WRKY38 (AT5G22570), WOX13 (AT4G35550) and DIV2 are potential LBD16 upstream regulators (**Fig. 3**). Among them, we show that *ZAT6* might be involved in regulating LR density, by binding to the *LBD16* promoter and upregulating *LBD16* expression in response to salt stress (**Fig. 4**). Although it is still unknown how ZAT6 is activated by salt in plant roots, ZAT6 was previously found to be expressed in the pericycle cells and to contribute to limiting Na^+^ accumulation in the shoots during salt stress^30^. ZAT6 was also reported to be phosphorylated directly by MITOGEN-ACTIVATED PROTEIN KINASE 6 (MPK6) and implicated in regulation of seed germination of Arabidopsis in saline conditions^31^. Salt may induce MPK6 activity within 15 min to activate downstream salt-responsive gene expression in Arabidopsis seedlings^32^. In addition, MPK3/MPK6 signaling is also involved in LR development via regulating downstream cell wall remodeling pathways^17, 33^. It is thus possible that salt might activate ZAT6 activity to promote LR formation via the MPK6-mediated signaling pathway.

Our cell wall monosaccharide composition analyses did not detect significant cell wall monosaccharide changes in the *zat6* mutant alleles in response to salt stress compared to Col-0 (**Fig. S10**), which may be explained by the fact that ZAT6 acts upstream to activate *LBD16* activity under salt to subsequently modulate the cell wall related genes. Nevertheless, salt significantly upregulated the expression of cell wall remodeling genes such as *PME2* and *XTH19* in Col-0, but not in the *35S::ZAT6::SRDX* line where both ZAT6 and LBD16 activities are suppressed (**Fig. 4d and Fig. S9),** suggesting that the ZAT6 -LBD16 module plays a central role in regulating cell wall remodeling genes under salt. Together with the similar pattern in the degree of pectin methylation between *zat6* alleles and the *lbd16-1* mutants (**Fig.5e**), our data support a role for *ZAT6* in cell wall remodeling, especially in affecting pectin methylation, via LBD16. It is likely that the misregulation of cell wall-related genes by salt application and consequent cell wall changes in the *zat6* and *lbd16-1* mutants compared to Col-0 may disturb the cell wall loosening processes during LR emergence, resulting in the reduced LR density in *zat6* and *lbd16* mutants compared to Col-0.

Although our results describe an auxin independent pathway controlling LR formation during long-term salt stress, auxin-ARF7/19-LAX3 signaling was also previously shown to induce expression of cell wall-related genes^16^, and auxin signaling in the endodermal cell layer is required to accommodate the swollen pericycle cells for LR initiation^34^. In addition, we showed that during early salt stress, the auxin-ARF7/19 module may still mediate LBD16 expression (**Fig. 2e**). We also found auxin responsive genes such as *PIN3*(*AT1G70940*)*, LAX1*(*AT5G01240*) and several *IAAs* (*IAA9*, *IAA11* and *IAA13*) are present in the upregulated GO term ‘response to hormone’ in our RNAseq analysis of *lbd16-1* mutants compared to Col-0 specifically after 6 h salt treatment, but not in control conditions (**supplementary dataset 4**), implying that there might be a feedback regulation of auxin signaling components during salt response through unknown mechanism(s). Recently, SUMOylation of ARF7 during hydropatterning was shown to block its binding activity to LBD16 and subsequently LR formation^2^. In addition, salt stress might activate auxin downstream TOR signaling which is required for LR formation by promoting the translation of ARF7/19 and LBD16^35, 36^. Although whether these above-mentioned post-transcriptional mechanisms could contribute to salt-induced root branching phenotype of *lbd16* mutants is still unknown, our results do provide evidence for a novel ZAT6-dependent transcriptional pathway to act independently of auxin in controlling LR development in salt.

In summary, our data show that salt inhibits root auxin levels and signaling and in parallel it activates alternative pathway(s) that help sustain root branching in salt. We have identified such a pathway -a salt-activated *ZAT6*-*LBD16* bypath - promoting root branching in response to salinity via altering cell wall remodeling. This provides a novel theoretical framework for developmental plasticity under stress; we propose that the identified ZAT6-LBD16 module allows plants to mitigate the effect of inhibition of the core developmental root branching program by salt, allowing optimization of root architecture in stressful conditions.

## Methods

### Plant materials, growth conditions and stress treatments

All the *Arabidopsis thaliana* wild-type, mutant and transgenic lines used in this research are in Columbia-0 (Col-0) ecotype background, except for the auxin reporter line C3PO (in Col-Utrecht ecotype)^23^. T-DNA knock-out insertion lines of *lbd16-1* (SALK_095791), *lbd16- 2* (SALK_040739) were obtained from the lab of Prof. Malcolm Bennett (University of Nottingham) and Prof. Ben Scheres (Wageningen University& Research), respectively. Both lines were genotyped again prior to being used in our study. T-DNA knock-out mutants of *zat6-1* (SALK_061991C), *zat6-2* (SALK_050196) were obtained from the Nottingham Arabidopsis Stock Centre (NASC, http://arabidopsis.info/) and genotyped for confirmation. The genomic complementation line genomic *LBD16::GFP* line in *lbd16-1* (gLBD16) and *pARF7::ARF7::GR* line was obtained from previously published report^13^. Primers used for genotyping are listed in **Supplementary Table 1**.

For seedling pregrowth in petri dishes for all the root phenotyping experiments, Arabidopsis seeds were surface sterilized in 70% (v/v) ethanol for 1 min, and then immersed in 30% (v/v) bleach with triton X-100 (10 µL/50 mL) for 10 minutes and followed by washing with sterilized MQ water for at least six times. Sterilized seeds were sown on 0.5x Murashige and Skoog (MS) (including B5 vitamins, Duchefa Biochemie B.V.) media containing 1% (w/v) 2-(N-morpholino) ethanesulfonic acid (MES) (Duchefa Biochemie B.V.) with PH=5.8 and 0.8% Plant Agar (Lab M, MC029), and then stratified at 4 °C in dark for two days followed by pre-growth at 21 °C under 16 h light/8 h dark cycle and 65-70% humidity.

For LR phenotype characterization, 4-day-old seedlings were transferred to the medium supplemented with 0 mM, 75 mM NaCl or 125 mM NaCl for 6 days.

For analyses of genomic *LBD16::GFP* expression patterns and *proLBD16::GUS* expression, 6-day-old seedlings were transferred into liquid 0.5x MS medium containing 1% (w/v) MES (Duchefa) (pH 5.8) with 0 mM or 125 mM NaCl on a shaker for 6 h to avoid hypoxia during the treatments before further microscopic analysis.

For time course expression studies of *LBD16* and *ZAT6,* Col-0 plants were grown hydroponically under 12 h/12 h light/dark cycle, 20°C until 4-week-old and then were treated with 0 mM or 125 mM NaCl for 0 h, 3 h, 6 h, 12 h, 18 h or 24 h.

For the RNAseq analysis, seeds were sown and pre-grown on 11 cm x 11 cm nylon mesh. Eight-day-old seedlings were transferred to 0.5x MS agar plates containing 1% (w/v) MES (pH 5.8) and 1% Dashin Agar with 0 mM or 130 mM NaCl for 6 h before roots were harvested for RNA isolation.

For cell wall analysis, seven-day-old seedlings of Col-0, *lbd16-1*, *zat6-1* and *zat6-2* grown on 11 cm x 1 cm nylon mesh strips were transferred to 0.5x MS agar plates containing 1% (w/v) MES (pH 5.8) and 0.8% Plant Agar with 0 mM or 125 mM NaCl for 24 h or 3 days (72 h) before roots were harvested for further analysis.

### Cloning and constructing

For yeast-one-hybrid assay, a 1309-bp *LBD16* promoter sequence upstream of the start codon was cloned into a gateway vector PDONR207 then was sequenced before the cassette containing LBD16 promoter was recombined into a modified gateway compatible destination bait vector pAbAi^37^ containing the AbAr gene (AUR-1C) reporter through LR reaction, then restriction enzyme digestions were performed to assess the final plasmid.

For *pLBD16::GUS* expression, *LBD16* promoter sequence was first cloned into PDONR207 then combined into pFAST-G04 binary vector for plant transformation and expression.

For *35S::ZAT6::SRDX* line, *ZAT6* coding sequence (with stop codon removed) was first cloned into PDONR221, then subcloned into pGWB119 vector for plant transformation.

For luciferase assay, PDONR221 carrying the 1309-bp *LBD16* promoter sequence was used for subcloning into pGWB435 carrying a LUC reporter; and PDONR221 carrying the *ZAT6* coding sequence was used for subcloning into pH7WG2 binary vector for overexpression.

All constructs for plant transformation were transformed into *Agrobacterium tumefaciens* AgL0 strain by electroporation and then transformed to Col-0 background by floral dipping.

### Root system architecture assay

Four-day-old seedlings of Col-0, *lbd16-1*, gLBD16 (in *lbd16-1*) complementation line, *zat6-1* and *zat6-2* mutants were transferred to 0.5x MS media containing 0 mM, 75 mM or125 mM NaCl in square petri dishes, and were then placed in 70-degree racks and scanned on day 6 (10-day-old seedlings) after transferring. Main root and LR phenotypes were traced by ImageJ with a Smartroot plugin^38^. Data were extracted in CSV files and then processed with R.

### Lateral root primordia assay

Four-day-old seedlings of Col-0, *lbd16-1* and *35S::ZAT6::SRDX* lines were transferred to 0.5x MS media containing 0 mM, 75 mM or 125 mM NaCl. Petri-dishes containing the transferred seedlings were placed in 70-degree rack at 21°C under 16 h light/8 h dark cycle and 65-70% humidity. For counting LRP and emerged LR under microscope, 6 days after transferring, seedlings were fixed by immersion first in 20% (v/v) methanol with 4% (v/v) hydrochloric acid at 57 °C for 20 min prior to immersion in 7% NaOH (w/v) in 60% (v/v) ethanol at room temperature for 15 min, followed by rehydration in 40%, 20% and 10% (v/v) ethanol for 5 min each. The fixed plants were stored in 10% ethanol at 4 °C prior to microscopy analysis. For microscopy analysis, Leica DM5200 microscope with Nomarski optics was used and the roots were immersed in 50% (v/v) glycerol on microscopy slides for scoring LRP. The LRP stages were determined according to previous report by Malamy and Benfey^5^.

### Yeast one hybrid

The final *pLBD16::AbAi* plasmid was transformed into pJ69-4α yeast strain as bait DNA, and the yeast library containing CDS of 1956 Arabidopsis transcription factors cloned into pDEST-22 vector was transformed into pJ69-4A yeast strain as prey proteins as previously described^39^. Positive yeast colonies were selected and confirmed by using different synthetic drop-out (SD) (Takara) media prepared according to the manufacturer’s instructions. Prior to screening, the strain carrying the *pLBD16::AbAi* bait was tested for autoactivation by using empty prey strains carrying empty vector of pDEST-22 under various concentrations of Aureobasidin A (AbA) (Takara) (0, 50 ng/mL,100 ng/mL, 150 ng/mL, 200 ng/mL and 400 ng/mL), and 200 ng/ml AbA was finally selected for further library screening. For mating-based screening, all the the strains carrying the prey library were grown overnight in SD-Trp medium in 96-well V bottom plates in a 28°C shaker. Yeast strain carrying the *pLBD16::AbAi* bait DNA was grown in a 28°C shaker overnight in selective medium (SD-Ura). Mating of yeast strains was carried out by spotting 5 µL 10-fold diluted overnight culture of both bait and prey strains in one spot on agar plate containing SD-glu complete at 28°C for 2∼3 days. The diploid yeast strain colony spots contained both *pLBD16::AbAi* bait and prey proteins were resuspend into 100 µL sterilized MQ and 5 µL was used to spot on selective agar media containing SD-Trp-Ura and 200 ng/mL AbA at 28 °C. Positive interactions were observed after 2 to 3 days.

### Histochemical GUS Staining

Six-days-old seedlings of *pLBD16::GUS* independent homozygous transformants in Col-0 backgrounds were transferred to 0.5x MS liquid media containing 0 mM or125 mM NaCl and placed on a shaker for 6 h during treatments. Then seedlings were transferred into GUS staining buffer containing 50 mM sodium phosphate buffer (pH 7.2), 10 mM EDTA, 0.1% (v/v) Triton X-100, 2.5 mM potassium ferri- and ferrocyanide, and 1 mg/mL 5-bromo4-chloro-3- indolyl-b-glucuronic acid (Duchefa) and vacuumed for 15 min at room temperature before incubation at 37°C for 1.5 h. Samples were cleared in clearing solution containing water (30 mL): chloral hydrate (80g):glycerol (10 mL) and mounted on microscope slides for imaging. A Leica DM5200 microscope with Nomarski optics was used for imaging with 20x dry objective.

### Confocal microscopy and auxin reporter analysis

For the detection of genomic *LBD16::GFP* (in *lbd16-1* background), 6-day-old seedlings were transferred to 0.5x MS liquid media containing 0 mM or 125 mM NaCl on a shaker for 6 h to avoid hypoxia. After treatment, roots were fixed using 4% paraformaldehyde (PFA) in 1 x PBS buffer for 1 h at room temperature with gentle agitation. After fixation, roots were washed twice for 1 min in 1 x PBS buffer. Roots were then transferred and immersed in ClearSee solution containing 10%(w/v) xylitol, 15% (w/v) sodium deoxycholate, and 25% (w/v) urea at room temperature with gentle agitation overnight. Seedlings cell membranes were counterstained for 30 min in a 0.1% Calcofluor white solution in ClearSee solution. Cleared and stained roots were mounted on slides with ClearSee solution for confocal imaging.

For auxin response analysis during salt stress, seven-day-old Arabidopsis Col-0 seedlings carrying a triple color auxin reporter C3PO line^23^ were transferred to 0.5x MS liquid media containing 0 mM or 125 mM NaCl and placed on a shaker for 6 h. After treatment, roots were fixed in PFA and cleared overnight using ClearSee solution. Cleared roots were mounted on slides with ClearSee solution. Roots were visualized using a Leica TCS SP5II confocal laser scanning microscope with a 40× NA = 1.20 water-immersion objective. mTurqoise2 and Calcofluor white were excited using a 405 nm diode laser, mVenus and GFP were excited using a 488nm Argon-ion laser, tdTomato was excited using a 552 nm laser. Detection was configured as follows: Calcofluor white was detected at 410-430 nm; mTurqoise2 was detected at 468-495 nm; GFP was detected at 500-550 nm; Venus was detected at 524-540 nm; tdTomato was detected at 571–630 nm. Z-stacks were acquired with 2.0 μm intervals, with the pinhole set to 2.0 Airy Unit. Virtual ratio images between channels were generated using the FIJI plugin Calculator Plus (Fiji). Ratios between pixel signal intensities of nuclei were calculated on SUM-slice projection of DII (C1;Venus) and mDII (C2;TdTomato) after subtracting background signal (value=4). Nuclei in each LRP area were manually selected as ‘Region of interest’ (ROI, Fiji) for quantifying average auxin input through ratio image (max C2/ max C1/ ROI) and average auxin output per ROI (max C3 / ROI).

### Luciferase assay

Dual Luciferase Reporter (DLR) system (Promega) was employed to test the activation of *pLBD16::LUC* by *35S::ZAT6*. Agrobacterium strains carrying *pLBD16::LUC*, *P19* gene-silencing repressor and *35S::ZAT6* (using an empty vector containing 35S promoter as control) constructs were co-infiltrated into leaves of four-week-old *Nicotiana Benthamiana*. *35S::Renilla-luciferase* (REN) was used as an internal control for transformation efficiency. Three days after infiltration, leaf samples of equal size were collected for protein extractions using the Dual-Luciferase Reporter Assay System kit (Promega, Fitchburg, USA). Glomax 96 microplate luminometer (Promega, Fitchburg, USA) was used to measure the luciferase activities. Using values from REN as internal control, the ratios were calculated.

### Public transcriptomic data analysis

For the analysis of public microarray datasets of LR development^26^ and root response to salt stress^25^, data analysis was performed using R with R packages provided by Bioconductor^40^. The R packages affy and simpleaffy were employed for reading .CEL files of the corresponding dataset, quality control, background adjustment and normalization^41, 42^. Quality control was performed by checking whether the scale factors of different chips were within 3-fold of one another and 3’/5’ ratios of each sample were below 3 using simpelaffy^42^. GCRMA was used to perform global normalization^43^. Principal component analysis was applied to check if the overall variability of the samples reflects their grouping. Based on the affy_ATH1_array_elements-2010-12-20 table in TAIR, probe sets that were annotated to hybridize to multiple loci in the Arabidopsis genome were removed from further analysis. Differentially expressed genes were determined using the LIMMA package in R^44^. The Benjamini-Hochberg method was used to correct for multiple testing. Genes with p-value ≤ 0.001 and at least two-fold expression difference between contrasts were considered as significantly differentially expressed genes (DEGs).

### Network inference

Network inference was performed using GRNBoost2^27^. The overlapping genes from public datasets and potential upstream regulators of *LBD16* from Y1H and published known upstream regulator(s) were used as input for the network inference^14^. The expression data for network inference were extracted from Geng et.al 2013^25^ to predict the *LBD16* upstream regulators under both control and salt conditions. Expression data from two timepoints for control treatment and six timepoints for salt conditions were used for the network prediction. To enhance the robustness of the network analysis, the algorithm was applied 100 times for expression data under different conditions (control and salt conditions). For each run, predicted direct regulators of *LBD16* were extracted from all inferred interactions using a custom Python script. The frequency of each predicted *LBD16*-related interaction was calculated over these 100 times of prediction. To prioritize the list of potential regulators of *LBD16*, only interactions with a frequency equal to or higher than 80% were listed for further analysis for the salt conditions, and interactions with frequency ≥ 40% were selected under control conditions to deal with higher variability in the network predictions (**Supplementary dataset 3**).

### RNAseq analysis

8-day-old seedlings were treated with 0 mM or 130 mM NaCl for 6 h. Roots were dissected for harvest and stored into liquid N2 for RNA isolation. Approximately 40 roots were pooled into one biological replicate. Total RNA was extracted using combined TRipure and RNA isolation kit (Qiagen). Three biological replicates were used for sequencing.

Total RNA was used for RNA library preparation suitable for Illumina HiSeq paired end sequencing using Illumina’s TruSeq stranded mRNA kit for polyA mRNA selection. Then mRNA was processed directly including RNA fragmentation, first and second strand cDNA synthesis, adapter ligation and final library amplification according to the manufacturer’s protocol. The final library was eluted in 30 µL elution buffer followed by quality assessment using a Bioanalyzer 2100 DNA1000 chip (Agilent Technologies) and quantified on a Qubit quantitation platform (Life Technologies).

Prepared libraries were pooled and diluted to 6 pM for TruSeq Paired End v4 DNA clustering on one single flow cell lane using a cBot device (Illumina). Final sequencing was carried out on an Illumina HiSeq 2500 platform using 126, 7, 126 flow cycles for sequencing paired end reads plus indexes reads. All steps for clustering and subsequent sequencing were done according to the manufacturer’s protocol. Reads were split per sample by using CASAVA 1.8 software (Illumina Inc, San Diego CA, USA). All sample preparations and sequencing were done by the Genomics lab of Wageningen University and Research, Business Unit Bioscience.

Read quality was assessed using FastQC (https://www.bioinformatics.babraham.ac.uk/projects/fastqc/) and low-quality reads were removed using “trim galore!” (https://www.bioinformatics.babraham.ac.uk/projects/trim_galore/). Reads were mapped to the Arabidopsis TAIR10 transcriptome using Salmon^45^.

Differential gene expression was quantified using DESeq2^46^ under different contrasts (*lbd16-1* vs Col0 in both control and salt conditions) using a false discovery rate (FDR) cut-off of 0.05. The counts of the top 5000 differentially expressed genes (DEGs) were used for principal component analysis. Expression heatmap of sample-to-sample distances was calculated from the Log2 transformed count data for overall gene expression. Clustering of 12 RNA-seq samples was carried out to show the overview of relationships between genotypes. GO enrichment analysis was performed for the NaCl-induced DEGs in *lbd16-1* vs Col-0 by using the R package gProfiler^47^.

### Gene expression study

For expression studies of *LBD16* in Col-0, *pARF7::ARF7::GR* in *arf7-1;arf19-1* double mutant line, *zat6-1*, *zat6-2,* and *35S::ZAT6::SRDX* line, 7-day-old seedlings pregrown on mesh strips were transferred to 0.5x MS agar plates containing 0 mM or 125 mM NaCl. The root samples were collected after 3 h, 6 h or 24 h treatment as specified in the figure legends for further analysis. For *LBD16* expression in *arf7-1;arf19-3* line, 5-day-old seedlings grown on mesh were transferred to 0 mM or 130 mM NaCl before roots were harvested for RNA extraction.

For time course gene expression study of *LBD16* and *ZAT6*, roots of 4-week-old hydroponically grown Col-0 plants were harvested after treated with 0 mM or 125 mM NaCl for 0 h, 3 h, 6 h, 12 h, 18 h and 24 h for RNA isolation.

For expression studies on *pARF7::ARF7::GR* in *arf7-1;arf19-1* double mutant lines, 7- day-old seedlings pregrown on mesh strips were transferred to 0.5x MS agar plates containing 0 mM or 125 mM NaCl for 3 h or 6 h before roots were harvested for RNA isolation.

Total RNAs were extracted with RNA isolation kits (NZYtech, Portugal). The RNA quality and concentration were assessed prior to cDNA synthesis. iScript cDNA synthesis kit (Bio-Rad) was used for cDNA synthesis. The primers used in qRT-PCR are listed in **Supplementary table 1.** 2x qPCRBIO SyGreen Mix Lo-ROX (PCRBIOSYSTEMS, UK) was used for qRT-PCR analysis. qRT-PCR analyses were performed using the CFX96 or CFX384 connect real-time PCR detection system (Bio-Rad).

### IAA measurement

Seven-day old seedlings were transferred to fresh 0.5x MS agar plates supplemented with 0 mM or 125 mM NaCl (Duchefa). After 1, 3, 6, 12 h being transferred, roots were dissected with a sharp blade, weighed, flash frozen in liquid N2 and stored at −80 °C for further analysis. For IAA extraction, approximately 60 roots (11.8±3.0 mg) were pooled in one biological replicate. Frozen material was ground to a fine powder and further extracted with 1 mL of 10% methanol containing 100 nM ^13^C-labeled IAA internal standard. The extraction was carried out according to a previous report with minor modification^48^. Namely, a StrataX 30 mg/3 mL spe-column (Phenomenex) was used.

For detection and quantification of IAA by liquid chromatography-tandem mass spectroscopy, sample residues were dissolved in 100 µL acetonitrile /water (20:80 v/v) and filtered using a 0.2 µm nylon centrifuge spin filter (BGB Analytik). IAA was identified and quantified by comparing retention time and mass transitions with IAA standard using a Waters XevoTQS mass spectrometer equipped with an electrospray ionization source coupled to an Acquity UPLC system (Waters) as previously described^49^. Chromatographic separations were conducted using acetonitrile/water (containing 0.1% formic acid) on a Acquity UPLC BEH C18 column (2.1 mm x100mm, 1.7 µm, Waters) at 40°C with a flowrate of 0.25 mL/min. The column was equilibrated for 30 min with acetonitrile /water (20:80 v/v) containing 0.1% formic acid. Samples were analyzed by injecting 5 µL, followed by the elution using program of 17 min in which the acetonitrile fraction linearly increased from 20% (v/v) to 70% (v/v). The column was washed after every sample by increasing the acetonitrile fraction to 100% in one minute and maintaining this concentration for 1 min. The acetonitrile fraction was reduced to 20% in one minute and maintained at this concentration for one minute before injecting the next sample. The capillary voltage was set at 3.5 kV, the source temperature and the desolvation temperature at 150 °C and 350 °C, respectively. Multiple reaction monitoring (MRM) was used for identification and quantification by comparing retention times and MRM transitions (+176.25>103.2; +176.25>130.2) with IAA standard. MRM transitions and cone voltages (30v) were set using the IntelliStart MS Console. Data acquisition and analysis were carried out using MassLynx 4.1 (TargetLynx) software (Waters) and was further processed in excel. The final IAA content in each sample was normalized by the corresponding internal standard recovery and sample weight. All values were normalized to the 1 h control timepoint.

### Cell wall analysis

Seven-day-old seedlings of Col-0, *zat6-1*, *zat6-2* and *lbd16-1* were transferred to 0.5x MS agar plates containing 0 mM or 125 mM NaCl. Roots were harvested into 2 mL tubes after 24 h or 72 h treatment with 0 mM or 125 mM NaCl and ground homogenously. 1 mL of pre-warmed 70% (v/v) ethanol was added to the tubes followed by vortexing. This solution was spun at max speed for 30 seconds and the supernatant discarded, after which this ethanol washing step was repeated and the supernatant was discarded again. A mixture of 50% (v/v) chloroform and 50% (v/v) methanol was added to the pellet, and the tube mixed by gently by inverting the tube at least 5 times. Tubes were spun at max speed for 30 seconds, and the supernatant was discarded. Acetone was added to the tubes, the tubes spun at max speed for 30 seconds, and the supernatant was discarded. This acetone wash was repeated 3 times. Tubes are left to dry with open lids overnight in a fume hood. The resulting alcohol insoluble residues (AIR) were used for further analysis.

For analysis of cell wall monosaccharide composition, 1-2 mg AIR was weighed out in 2 mL screw caps tubes and used for extraction of neutral cell wall sugars and uronic acids as described^50^.High-performance anion-exchange chromatography with pulsed amperometric detection (HPAEC-PAD) was performed on a biocompatible Knauer Azura HPLC system, equipped with an Antec Decade Elite SenCell detector. Monosaccharides were separated on a Thermo Fisher Dionex CarboPac PA20 column BioLC guard (3 x 30 mm) and analytical (3 x 150 mm) column as previously described (https://doi.org/10.1073/pnas.2119258119). Briefly, samples were diluted with ultrapure water, ribose was added as an internal standard and separation performed with a solvent gradient of (A) water, (B) 10 mM NaOH and (C) 700 mM NaOH at 0.4 mL /min flow rate. 0 to 25 min: 20% B, 25 to 28 min: 20 to 0% B, 0 to 70% C, 28 to 33 min: 70 % C, 33 to 35 min: 70 to 100% C, 35 to 38 min: 100% C, 38 to 42 min: 0 to 20% B, 100 to 0% C, 42 to 60 min: 20% B.

The degree of pectin methyl-esterification was analyzed as described^51^ with minor modifications. 1 mg AIR was saponified in 0.2 mL 250 mM NaOH for 1h at room temperature and neutralized with HCl. After centrifugation, 50 µL diluted supernatant was incubated with an equal volume of 0.1 M sodium phosphate buffer (pH 7.5) containing 0.03 units alcohol oxidase (Sigma A2404) for 15 min at room temperature under shaking. After addition of 100 µL 0.02 M 2,4-pentanedione in 2 M ammonium acetate and 0.5 M acetic acid, samples were incubated at 68°C for 10 min, briefly cooled on ice and transferred in 96-well microtiter plates. Absorbance was measured at 412 nm in a Tecan infinite 200Pro microplate reader and calibrated against a 0-350 µM formaldehyde standard series. The degree of pectin methyl-esterification was calculated as the molar ratio of methanol to uronic acid determined via HPAEC-PAD analysis.

### Statistical analysis

Graph Pad Prism software and R were used for statistical analysis. For the proportion of non-emerged LRP and emerged LRs a generalized linear model was fitted (glm(cbind(‘non-emerged’,emerged) ∼ genotype*condition, family = binomial(link = “logit”)) and statistical differences were derived from least squares mean analysis with custom contrast (emmeans R package 1.8.1-1).

## Author contributions

Y.Z. and C.T. conceived the project. Y.Z. performed most of the experiments and the data analysis. Y.L. performed the network inference under the supervision of A.D.J.D and Y.Z. Y.L conducted RNAseq data analysis, assisted by J.L.. T.Z. assisted with the experiments and performed confocal microscope analyses. K.D. contributed to the qRT-PCR analysis and phenotypic analysis of the *35S::ZAT6::SRDX* lines. D.K. performed the phenotypic analysis of the *lbd16-2* allele and assisted the statistical analysis for Fig.1c. J.L. and F.V. performed IAA analysis. K.M. and T.E. contributed the cell wall monosaccharide composition analysis. H.L., J.M., N.G-B, J.Y., Yu.Z. Y.W., T.G., and J.W assisted with experiments and data analysis. A.D.J.D supervised the network inference and contributed to the RNAseq data analysis. Y. Z and C.T. acquired the funding, guided the research and wrote the manuscript. All authors approved the manuscript.

## Acknowledgements

We would like to thank Malcolm Bennett (University of Nottingham) for kindly providing us the *gLBD16::GFP* (*lbd16-1*) line and *arf7-1;arf19-3* double mutant, Tom Beeckman (Ghent University) and Hidehiro Fukaki for sharing with us the *pARF7::ARF7::GR* (*arf7-1;arf19-1*) line, and Dolf Weijers (Laboratory of Biochemistry, Wageningen University & Research) for providing the C3PO line. We thank Elio Schijlen (Bioscience, Wageningen University & Research) for the assistance in library preparation and RNA sequencing. This work is supported by the European Research Council (ERC) (grant agreement 724321), Dutch Research Council (NWO) Vici grant (Grant VI.C.192.033) and NWO-ENW KLEIN (OCENW.KLEIN.421), Netherlands Organisation for Health Research and Development (ZonMW) Enabling Technologies Hotels programme-early career scientists projects (Grant No.435004012) and a China Scholarship Council (CSC) - Sino-Dutch Bilateral Exchange Scholarship (CF 13731).

