## Supplementary figures and table for "Root branching in salt requires auxin-independent modulation of LBD16 function"

**Extended data**

**Fig. S1** Root phenotypic analysis of *lbd16-2* mutants compared to Col-0 under control and salt conditions.

**Fig. S2** Root phenotypic analysis of Col-0, *lbd16-1* and genomic LBD16 complementation line (*gLBD16::GFP* in *lbd16-1* background).

**Fig. S3** *pLBD16::GUS* activity in main root zones and developing lateral root primordia (LRP) of Col-0 seedlings.

**Fig. S4** Salt-affected auxin response in the main root and early stages of lateral root primordia (LRP) of Arabidopsis wild-type plants and expression of *LBD16* in *pARF::ARF7::GR* (*arf7-1;arf19-1*).

**Fig. S5** Characterization of LBD16 upstream transcription factors in the root by yeast-one-hybrid screening followed by network inference and expression of ZAT6 in Col-0, *arf19-3; arf7-1* mutants and *pARF7::ARF7::GR* (*in arf7-1;arf19-1*) plants.

**Fig. S6** Characterization of T-DNA knockout alleles of ZAT6 and their root phenotypic identification.

**Fig. S7** Root phenotypic identification of *35S::ZAT::SRDX* line compared with Col-0.

**Fig. S8** Quality control of RNAseq analysis of Col-0 and *lbd16-1* mutant and identification of cell-wall related GO term.

**Fig. S9** Expression of *PME2* and *XTH19* in the roots of *35S::ZAT6::SRDX* line.

**Fig. S10** Cell wall analysis of wild-type Col-0 and loss-of-function mutants of *LBD16* and *ZAT6* in response to salt stress.

**Table S1** Primers used in this study.

**Fig. S1**

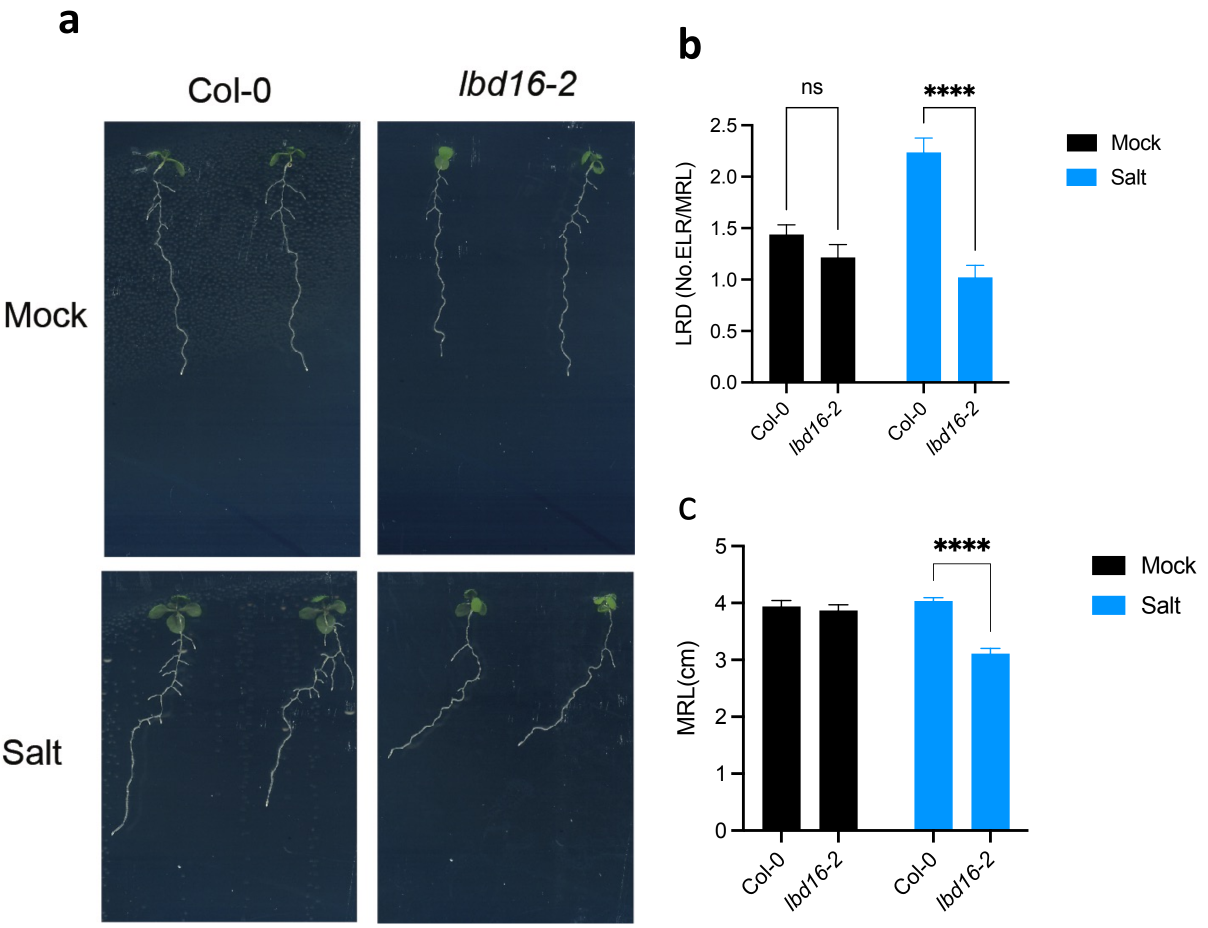


**Fig. S1 Root phenotypic analysis of *lbd16-2* mutants compared to Col-0 under control and salt conditions. a,** Representative images of *lbd1-2* compared with Col-0 under 0 mM, 75 mM NaCl. **b,** Emerged lateral root (ELR) density of *lbd16-2* compare to Col-0 under control (on 6-day old seedlings) and 75 mM NaCl condition (on 10-day old seedlings). Four-day-old seedlings were transferred to agar plates containing 0 mM NaCl or 75 mM NaCl with addition of 1% sucrose for 2 or 4 days before roots were scanned for emerged lateral root number quantification. **c,** Main root length of Col-0 and *lbd16-2* (n=20~25). Data in b and c represent means ± SEM. Statistical analyses in b and c were done using two-way ANOVA followed by Tukey's multiple comparison tests. **** P<0.001.

**Fig. S2**

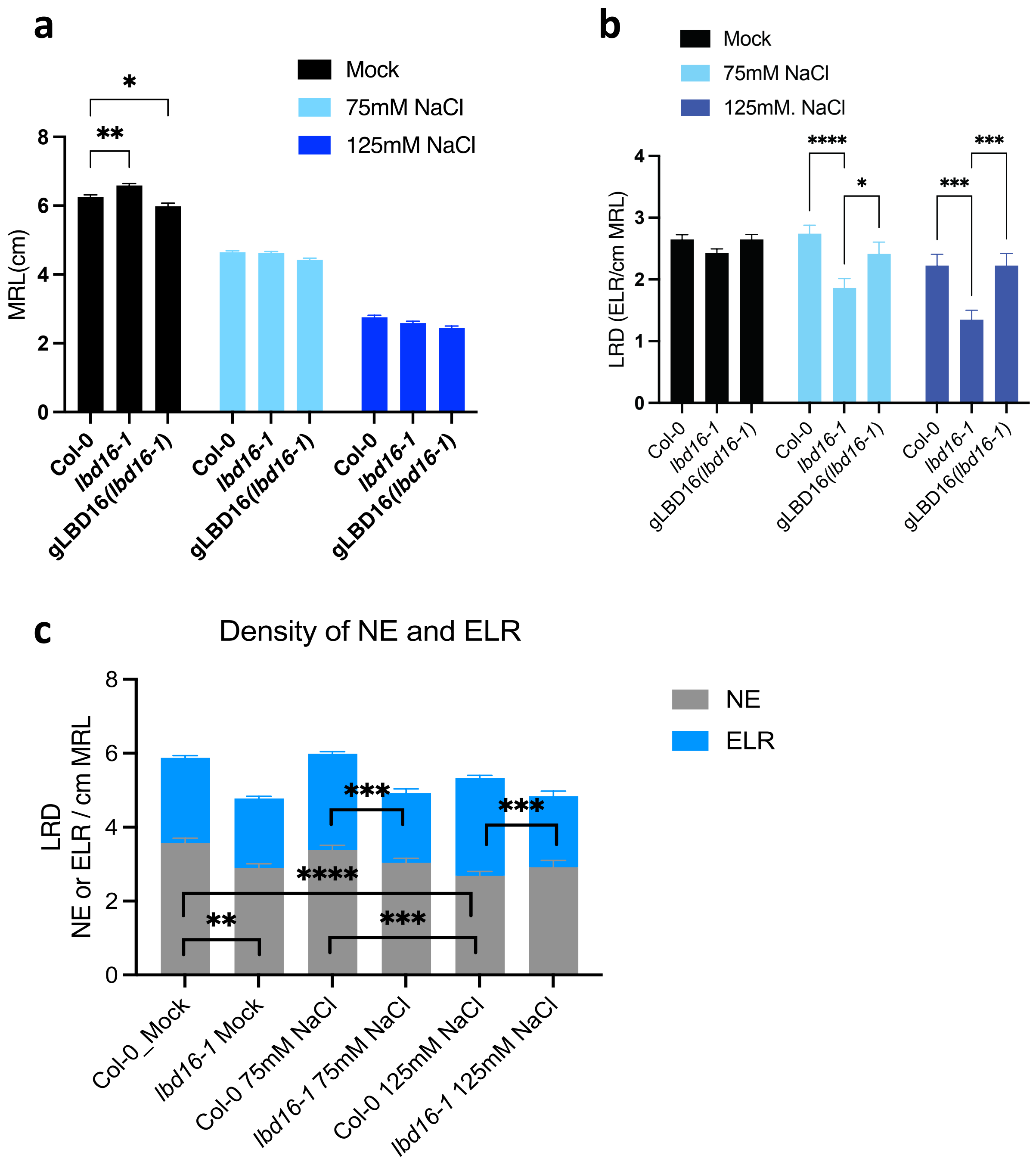


**Fig. S2 Root phenotypic analysis of Col-0, *lbd16-1* and genomic *LBD16* complementation line (*gLBD16::GFP* in *lbd16-1* background). a,** Main root length of Col-0, *lbd16-1* and genomic LBD16 complementation line (*gLBD16::GFP* in *lbd16-1* background) (n=20). Data are representative of at least 3 independent experiments. Statistical analysis was done using two-way ANOVA followed by Tukey's multiple comparison tests. * P<0.05, ** 0.05<P<0.01. **b,** Density of emerged lateral roots (ELR) in Col-0, *lbd16-1* mutant and gLBD16 (*lbd16-1*) complementation line in control and salt conditions (0 mM, 75 mM and 125 mM NaCl) (n= 15~20). **c,** Density of non-emerged lateral root primordia (NE) and ELR in *lbd16-1* compared with Col-0 under control and salt conditions (n=25~34, a pool of 3 independent experiments).

Four-day-old seedlings were transferred to agar plates containing 0 mM,75 mM or 125 mM NaCl for 6 days (10-day old seedlings) before roots were scanned for root clearing and quantification of ELR and NE under microscope. Data in a-c represent means ± SEM. Statistical analyses in a-c were done using two-way ANOVA followed by Tukey's multiple comparison tests. ** $0.05<P<0.01$., ***$0.01<P<0.001$, **** $P<0.001$.

**Fig. S3**

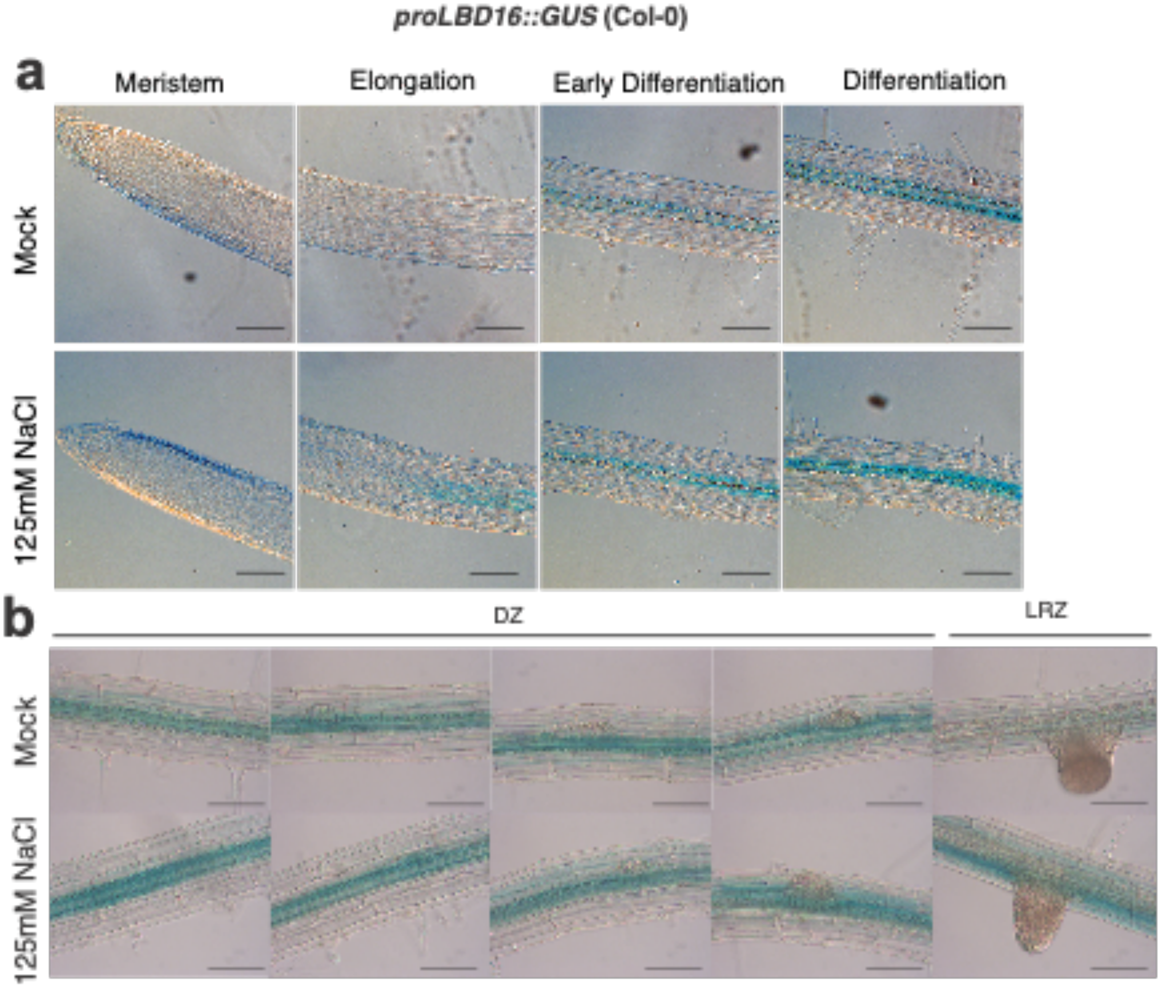


**Fig. S3 *pLBD16::GUS* activity in main root zones and developing lateral root primordia (LRP) of Col-0 seedlings. a**, *LBD16* promoter activity indicated by GUS staining in different root zones after treated with or without 125mM NaCl. **b,** *pLBD16::GUS* activity in developing LRP under control and 125 mM NaCl. Seven-day old seedings were treated with 0 mM or 125 mM NaCl for 6 h prior to GUS staining. Scale bars represent 0.1mm in a and b. DZ denotes differentiation zone; LRZ denotes lateral root zone (root zone with emerged LR).

**Fig. S4**

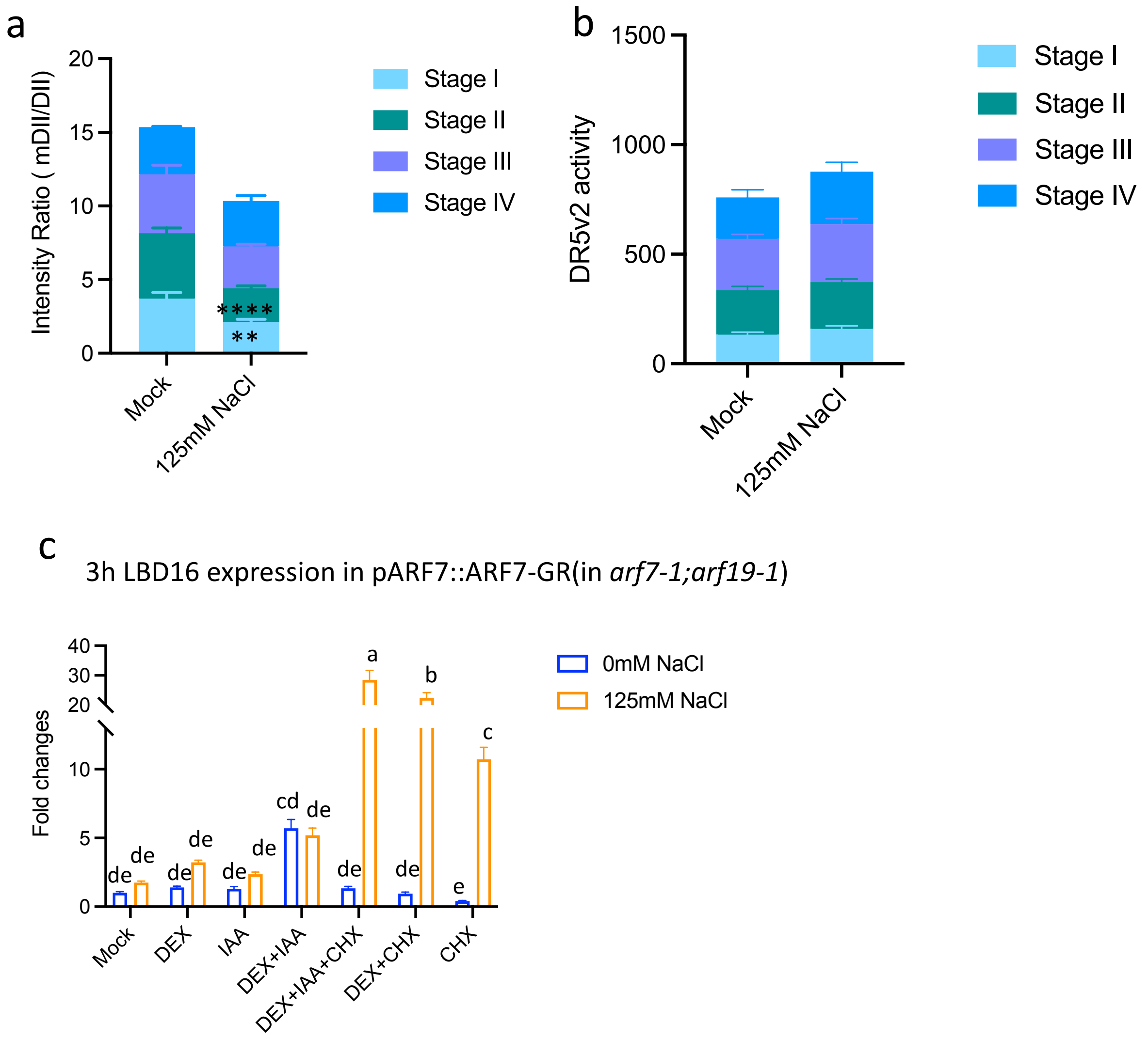


**Fig. S4 Salt-affected auxin response in the main root and early stages of lateral root primordia (LRP) of Arabidopsis wild-type plants and expression of *LBD16* in *pARF::ARF7::GR* (*arf7-1;arf19-1*). a,** auxin input signal indicated by the ratio of mDII/DII in developing LRP (stage I through IV) of 6-day old wild-type Col-Utrecht plants in control and salt (treated with 125 mM NaCl for 6 h) conditions. **b,** auxin output DR5v2 activity in early stages of LRP of 6-day old wild-type Col-Utrecht plants in control and salt (treated with 125 mM NaCl for 6 h) conditions. **c,** Relative expression of *LBD16* in the *pARF7::ARF7::GR* (*in arf7-1;arf19-1*) plants were treated with 1 µM IAA, 2 µM dexamethasone (DEX) and/or 10 µM cycloheximide (CHX) inducible lines in control and salt conditions after 3 h 125 mM NaCl treatment in comparison with mock treatment. Expression values were normalized by house-keeping gene At2g43770. Data in a- c represent means ± SEM. Statistical analyses in a and b were done using two-sided T-test. Statistical analysis in c was done using two-way ANOVA followed by Tukey's multiple comparison tests. ** 0.05<P<0.01, **** P < 0.001.

### Fig. S5

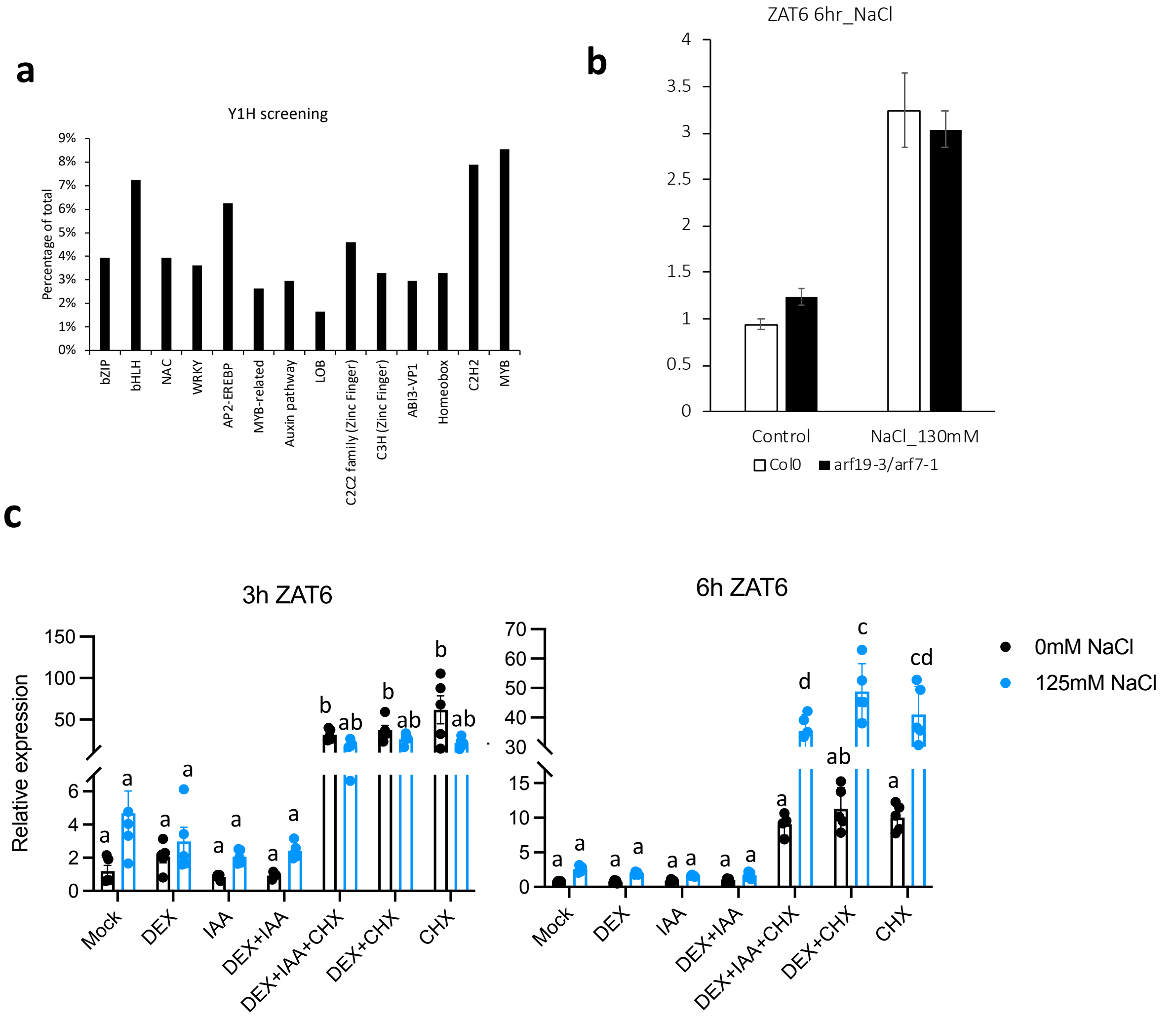


**Fig. S5 Characterization of LBD16 upstream transcription factors in the root by yeast-one-hybrid screening followed by network inference and expression of ZAT6 in Col-0, *arf19-3; arf7-1* mutants and *pARF7::ARF7::GR* (*in arf7-1;arf19-1*) plants. a**, Distribution of transcription factor families as putative upstream regulators of LBD16 via yeast-one-hybrid screening. **b,** Venn diagram showing the number of candidate genes that were identified as putative upstream regulators of *LBD16* under control and salt conditions (and overlaps between control and salt) from the network inference. **c,** Expression of *ZAT6* in *pARF7::ARF7::GR* (*in arf7-1;arf19-1*) plants treated with 1 µM IAA, 2 µM dexamethasone (DEX) and/or 10 µM cycloheximide (CHX) after 3 h or 6 h 125 mM NaCl treatment in comparison with mock treatment (n=4). **d**, Expression of *ZAT6* in Col-0 and *arf19-3; arf7-1* mutants in control and salt conditions (n=4~5). Expression values in c and d were normalized by house-keeping gene *At2g43770*. Data in c and d represent means ± SEM. Statistical analyses in c and d were done using two-way ANOVA followed by Tukey's multiple comparison tests and two-sided T-test, respectively.

**Fig. S6**

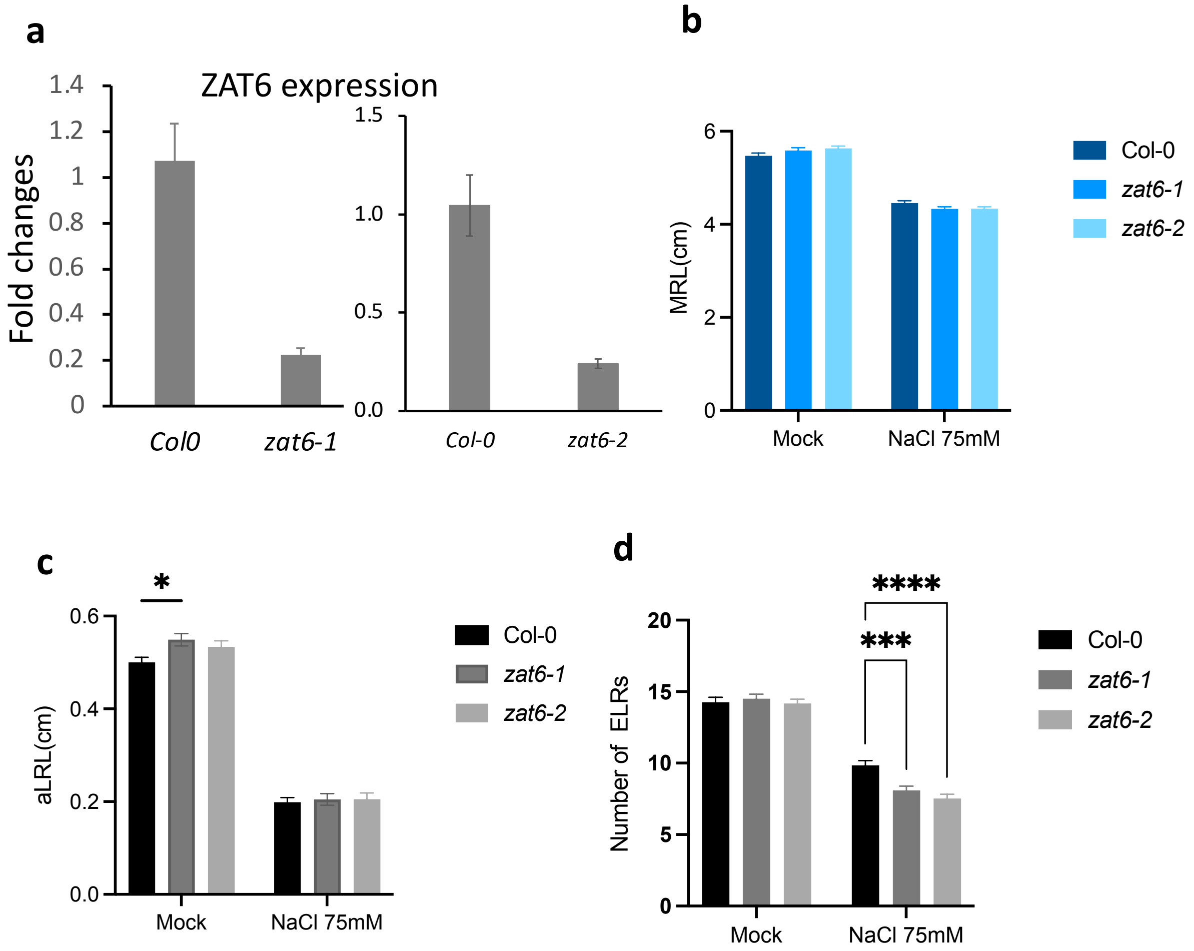


**Fig. S6 Characterization of T-DNA knockout alleles of ZAT6 and their root phenotypic identification. a**, qRT-PCR confirmation of the expression of *ZAT6* in the T-DNA knockout *zat6-1* and *zat6-2* lines (n=4). The expression is fold change normalized by house-keeping gene in comparison to Col-0. **b,** Main root length of 10-day old seedling of Col-0, *zat6-1* and *zat6-2*. **c,** Average lateral root length in 10-day old seedling of Col-0, *zat6-1* and *zat6-2* after transferring to 0 mM or 75 mM for 6 days. **d,** Number of emerged lateral roots (ELR) in 10-day old seedling of Col-0, *zat6-1* and *zat6-2* after transferring to 0 mM or 75 mM for 6 days. Data in a-d represent means ± SEM. Statistical analyses in a was done using two-sided T-test, and in b- d were done using two-way ANOVA followed by Tukey's multiple comparison tests. * $P<0.05$, ***$0.001<P<0.01$, **** $P < 0.001$.

#### Fig. S7

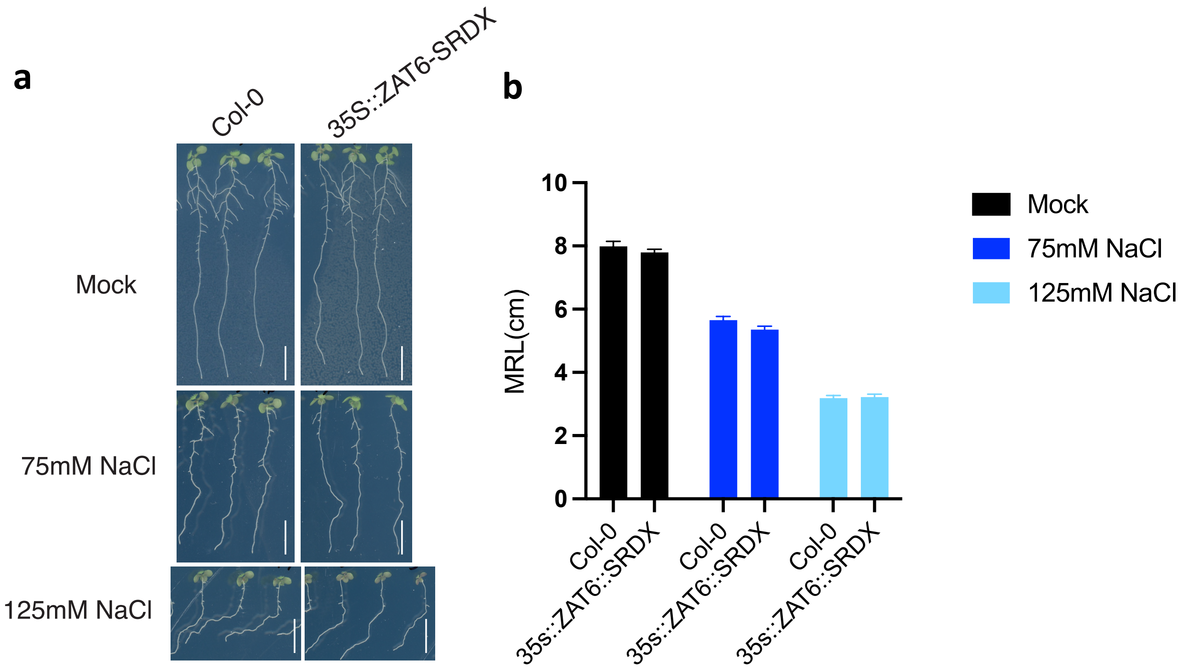


**Fig. S7 Root phenotypic identification of *35S::ZAT::SRDX* line compared with Col-0. a,** Representative images of root phenotypes of 10-day old seedlings of Col-0 and *35S::ZAT::SRDX* line under mock, 75 mM NaCl and 125 mM NaCl conditions. Scale bars represent 1cm. **b,** Main root length of 10-day old seedling of Col-0 and *35S::ZAT::SRDX* line. Data in b are representative of two independent experiments (n=10~15). Data in b represent means ± SEM. Statistical analyses in b was done using two-way ANOVA followed by Tukey's multiple comparison tests.

#### Fig. S8

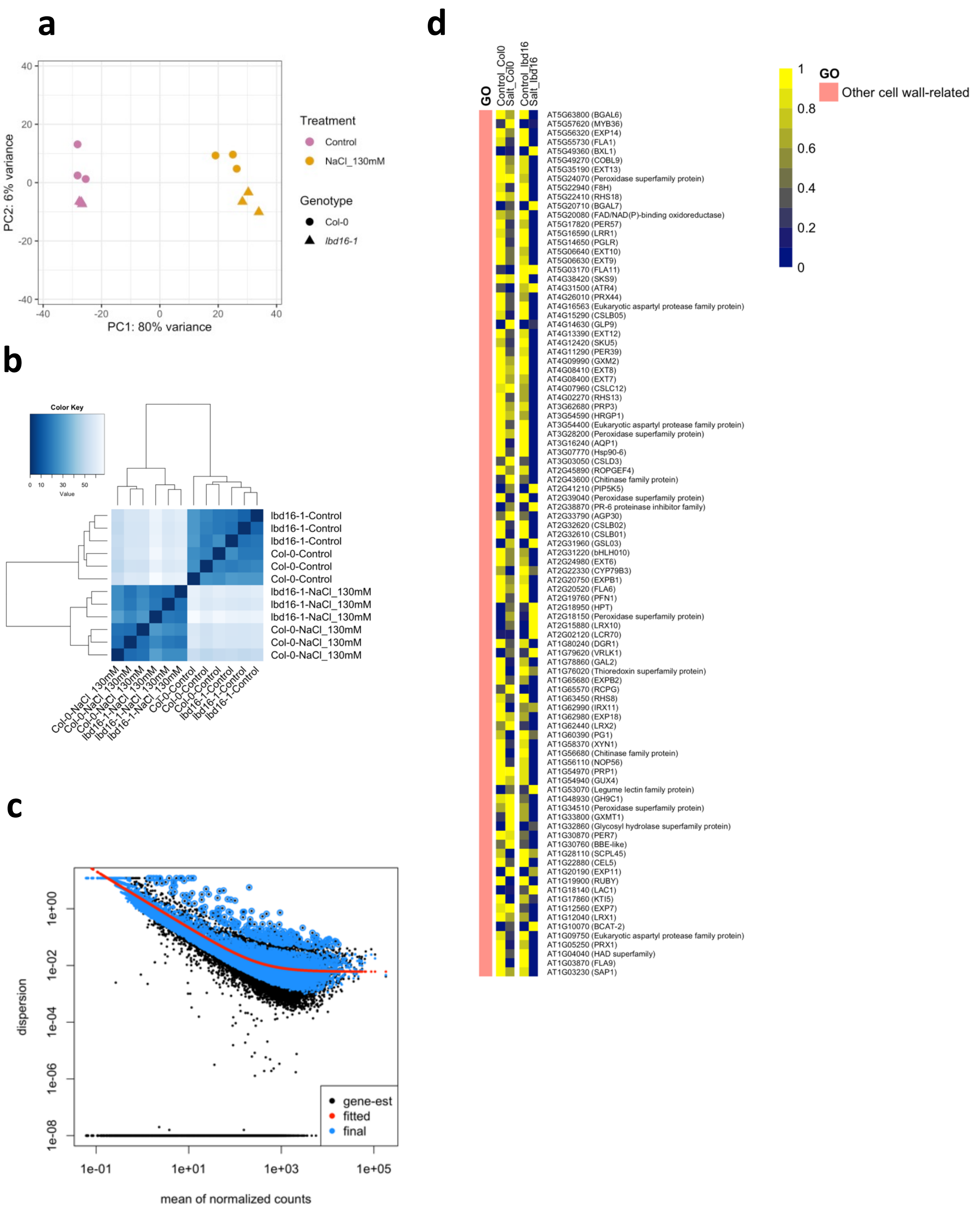


**Fig. S8 Quality control of RNAseq analysis of Col-0 and *lbd16-1* mutant and identification of cell-wall related GO term. a**, PCA of top 5000 differentially expressed genes in the RNA-seq analysis. Each dot represents a sample. Color coded according to the treatments, and shape of dot coded according to the time points. Samples with similar gene expression profiles are clustered together, which provides an indication of the reproducibility of the biological replicates and the quality of the sequencing. **b**, Clustering of the samples in the RNAseq analysis. The Euclidean distance of the log2-transformed counts were shown in the heatmap. A darker color in the heatmap means the corresponding samples are more correlated. **c**, A dispersion plot of the RNA-seq analysis processed by Deseq2. The dispersion decreases smoothly for genes with higher expression and eventually reaches an asymptote, which can be considered as the biological variability that is present in the dataset. **d,** Expression profiles of genes from other cell wall -related GO term (besides pectin and xyloglucan-related GO terms shown in Fig.5) in Col-0 and *lbd16-1* under control and salt conditions.

#### Fig. S9

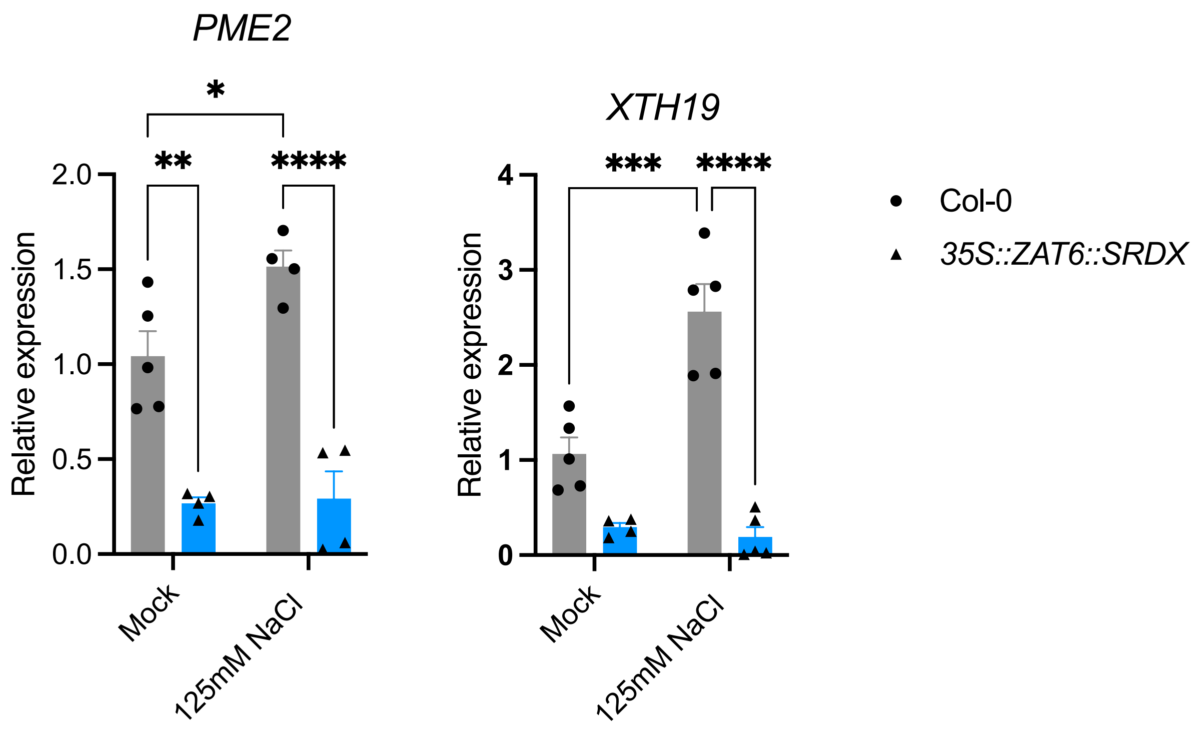


**Fig. S9 Expression of *PME2* and *XTH19* in the roots of *35S::ZAT6::SRDX* line.** The expression is fold change normalized by house-keeping gene (*At2g43770*) in comparison with Col-0 under mock conditions (n=4~5 pools of 40~45 roots). Roots of 7-day old seedlings of Col-0 and *35S::ZAT6::SRDX* line were treated with 0 mM or 125 mM NaCl for 24 h before they were harvested for gene expression analysis. Data represent means ± SEM. Statistical analyses were done using two-way ANOVA followed by Tukey's multiple comparison tests. * P <0.05, ** 0.05<P<0.01, *** P<0.01, **** P<0.001.

#### Fig. S10

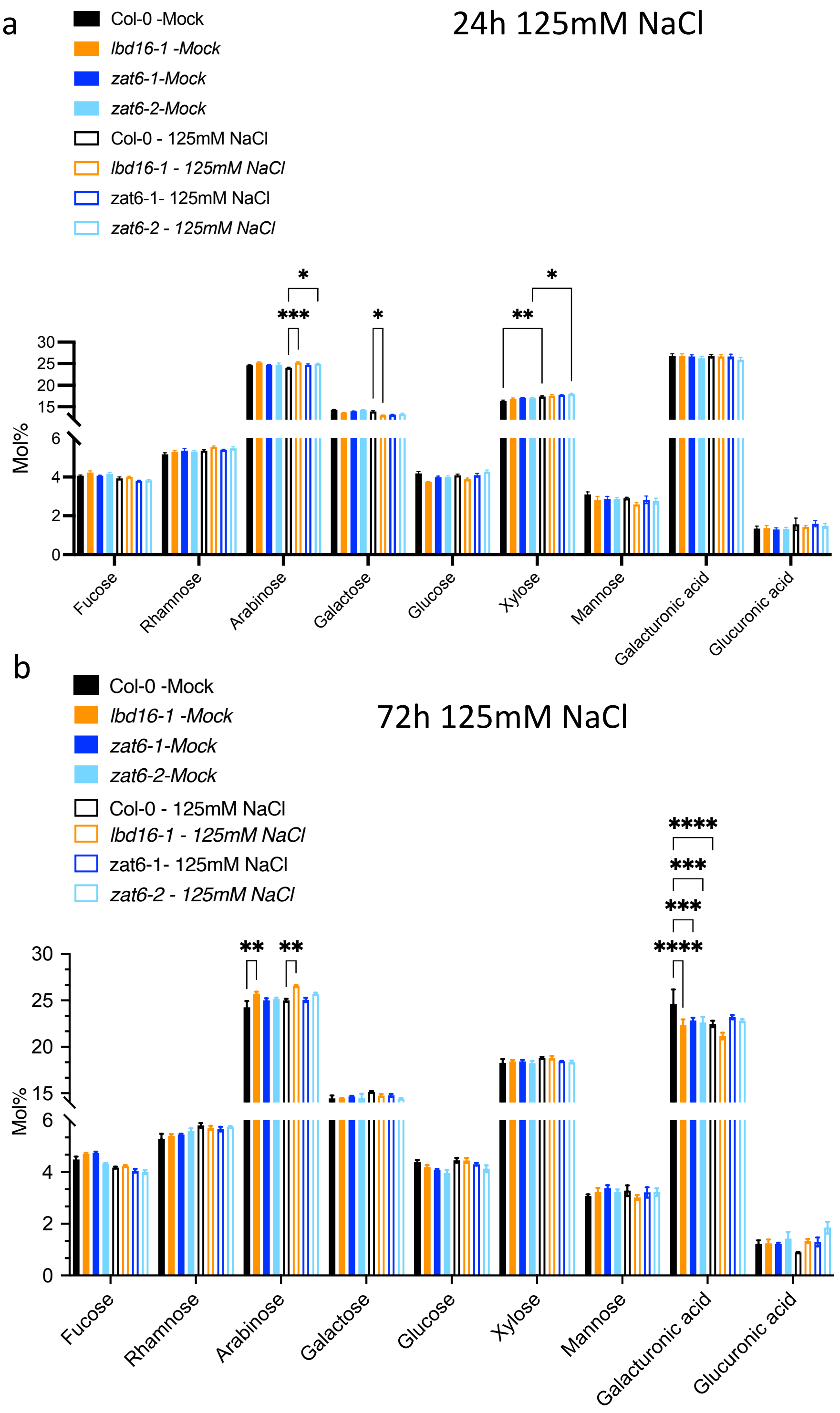


**Fig. S10 Cell wall analysis of wild-type Col-0 and loss-of-function mutants of *LBD16* and *ZAT6* in response to salt stress. a**, Cell wall monosaccharides composition analysis in roots of 7-day old Col-0, *lbd16-1*, *zat6-1* and *zat6-2* mutants after 24 h treatment with 0mM or 125mM NaCl (n=3~4). **b**, Cell wall monosaccharide composition analysis in roots of 7-day old Col-0, *lbd16-1*, *zat6-1* and *zat6-2* mutants after 72 h treatment with 0 mM or 125 mM NaCl (n=3~4). Data in a and b were obtained from 3~4 pools of samples containing 60~80 roots. Data in a and b represent means ± SEM. Statistical analyses in **a** and **b** were done using two-way ANOVA followed by Tukey's multiple comparison tests. * P<0.05, ** 0.01<P<0.05, ***0.001<P<0.01, **** P < 0.001.

**Supplementary Table S1** Primers used in this study.

| Primer name | Sequence (5'-3') | Additional information |
|---|---|---|
| Geno_*lbd16-1*_F | TTTCTTCCTTTTTGCTTTGCC | SALK_095791 genotyping gene-specific forward primer |
| Geno_*lbd16-1*_R | CAATGGCCAGTGACTTAAAGC | SALK_095791 genotyping gene-specific reverse primer |
| Geno_*lbd16-2*_F | CGCCAAAATCTTGAGTAAACG | SALK_040739 genotyping gene-specific forward primer |
| Geno_*lbd16-2*_R | TCATTTCTGTTTCAATTCTCCG | SALK_040739 genotyping gene-specific reverse primer |
| Geno_*zat6-1*_F | AGTAAGCGAAAAGCTTTTCCG | SALK_061991C genotyping gene-specific forward primer |
| Geno_*zat6-1*_R | GGGGCACTATAGTGGCACTAG | SALK_061991C genotyping gene-specific reverse primer |
| Geno_*zat6-2*_F | AGACGAAGAAGAAGGCAGGTC | SALK_050196 genotyping gene-specific forward primer |
| Geno_*zat6-2*_R | TCGTACTTTGGCGAAACATTC | SALK_050196 genotyping gene-specific reverse primer |
| Left border primer SALK T-DNA genotyping | ATTTTGCCGATTTCGGAAC | SALK left border primer for genotyping |
| qP_LBD16_F | TCATCATCAAACCGGAGGAG | AT2G42430 qPCR forward primer |
| qP_LBD16_R | GCCTGAAGCTCACCTAAATCG | AT2G42430 qPCR reverse primer |
| qP_ZAT6_F | TACCGGAATTCTCGATGGTC | AT5G04340 qPCR forward primer |
| qP_ZAT6_R | GAAGAAGAACATCAATCAAAATCG | AT5G04340 qPCR reverse primer |
| qP_PME2_F | GACGGAAGCGGTGACTTTAC | AT1G53830 qPCR forward primer |
| qP_PME2_R | ATAGTTTTGCCACGGCCATC | AT1G53830 qPCR reverse primer |

| | | |
|---|---|---|
| qP_XTH19_F | CTTGTAGCCCAATGCTCTGC | AT4G30290 qPCR forward primer |
| qP_XTH19_R | GAAACTTGTCCCTGGTAACTCTG | AT4G30290 qPCR reverse primer |
| at2g43770 | TATCATTGGATCTTGCAGTAGTG | Housekeeping gene, Dekkers et al., 2012 |
| at2g43770 | ACATCGTCGATTCTAAAGACTTC | Housekeeping gene, Dekkers et al., 2012 |
| *ZAT6*_CDS_P1P2_F | ggggacaagtttgtacaaaaaagcaggctacATGGCACTTGAAACTCTTACTTCTC | AT5G04340 For gateway cloning into pDONR221/207 |
| *ZAT6*_CDS_P1P2_R | ggggaccactttgtacaagaaagctgggttGGGTTTCTCCGGGAAGTC | AT5G04340 For gateway cloning into pDONR221/207 |
| Pro_pLBD16_F | gcggaagaacttataaaataacttttctaaaattaaaat | LBD16 promoter cloning |
| Pro_pLBD16_R | cggcgaaacgaacaaaaaag | LBD16 promoter cloning |
| P1P2_pLBD16_F | GGGG ACA AGT TTG TAC AAA AAA GCA GGC TAC gcggaagaacttataaaataacttttctaaaattaaaat | LBD16 promoter gateway cloning |
| P1P2_pLBD16_R | GGGG AC CAC TTT GTA CAA GAA AGC TGG GTT cggcgaaacgaacaaaaaag | LBD16 promoter gateway cloning |
